# A whole-genome-based approach for estimation and characterization of individual inbreeding

**DOI:** 10.1101/106765

**Authors:** T. Druet, M. Gautier

**Author notes:** Corresponding author: Tom Druet, Unit of Animal Genomics, GIGA (B34 +1), Quartier Hôpital, Avenue de l’Hôpital, 11, B-4000 Liège, Belgium.

## Abstract

Inbreeding results from the mating of related individuals and has negative consequence because it brings together deleterious variants in one individual. Inbreeding is associated with recessive diseases and reduced production or fitness. In general, inbreeding is estimated with respect to a base population that needs to be defined. Ancestors in generations anterior to the base population are considered unrelated. We herein propose a model that estimates inbreeding relative to multiple age-based classes. Each inbreeding distribution is associated to a different time in the past: recent inbreeding generating longer homozygous stretches than more ancient. Our model is a mixture of exponential distribution implemented in a hidden Markov model framework that uses marker allele frequencies, genetic distances, genotyping error rates and the sequences of observed genotypes. Based on simulations studies, we show that the inbreeding coefficients and the age of inbreeding are correctly estimated. Mean absolute errors of estimators are low, the efficiency depending on the available information. When several inbreeding classes are simulated, the model captures them if their ages are sufficiently different. Genotyping errors or low-fold sequencing data are easily accommodated in the hidden Markov model framework. Application to real data sets illustrate that the method can reveal recent different demographic histories among populations, some of them presenting very recent bottlenecks or founder effects. The method also clearly identifies individuals resulting from extreme consanguineous matings.

## Introduction

With his pioneering work on self-fertilization, Darwin early noticed that mating relatives generally leads to off-spring with a reduced fitness (Darwin, 1876). This phenomenon now referred to as inbreeding depression may mostly result from an increased homozygosity for (recessive) deleterious variants although a lack of heterozygosity at loci displaying heterozygous advantage (overdominance) might also be involved (Charlesworth & Willis, 2009). Accordingly, populations displaying high levels of individual inbreeding show a higher prevalence of monogenic disorders (e.g., Charlier *et al*, 2008) or complex diseases (e.g., Rudan *et al*, 2003). Inbreeding depression can thus increase the risk of extinction by reducing the population growth rate (Hedrick & Kalinowski, 2000; Keller & Waller, 2002) although it may be conversely favorable in some conditions by purging deleterious variants from the population (Estoup *et al*, 2016). Assessing individual inbreeding is then of paramount interest to improve the management of populations under conservation or selection, and from a more general evolutionary perspective to better understand the genetic architecture of inbreeding depression.

The first standard measure for the level of individual inbreeding was introduced by Wright (1922) as the coefficient of inbreeding (*F*) that he defined in terms of correlations between the parents uniting gametes. Further, Malécot (1948) proposed an alternative and more intuitive probabilistic interpretation of *F* as the probability that any two genes each randomly sampled in the parents gametes are identical by descent (IBD), i.e., are themselves derived from a common ancestor. In practice, estimation of *F* has long been only feasible using pedigree data and was hence limited to a few populations where such information had been recorded. Nevertheless, pedigrees remain usually limited to a few past generations leading to downward bias in the estimates of *F* since remote relationships are ignored (Keller *et al*, 2011), and they might also contain a non negligible proportion of errors even in well recorded domestic breeds (Leroy *et al*, 2012). In addition, whatever the pedigree depth and accuracy, pedigree-based estimates of *F* are only providing the expected proportion of individual genomic inbreeding which might departs from the actual genomic inbreeding due to mendelian sampling and linkage (Hill & Weir, 2011). With the advent of next generation sequencing and genotyping technologies, using genomic information to estimate the (realized) individual inbreeding proved particularly valuable (Wang, 2016) opening new avenues in the study of inbreeding in a wider range of populations including wild ones since genealogy is no more required (Hedrick & Garcia-Dorado, 2016; Kardos *et al*, 2016).

Genomic approaches to estimate *F* basically rely on the identity by state (IBS) status of genotyped markers and may be divided in two broad categories depending on whether or not they use linkage map information. The first type of methods ranges from simple estimates of individual heterozygosities (e.g., Szulkin *et al*, 2010) or homozygosities (e.g., Bjelland *et al*, 2013) to more advanced approaches based on the estimation of the realized genomic relationship matrix (VanRaden, 2008; Yang *et al*, 2010) or moment-based estimators to correct for population-structure in the estimation of population allele frequencies (e.g., Manichaikul *et al*, 2010). Their accuracy depends strongly on the number and informativeness of the genotyped markers (Kardos *et al*, 2015) but they always remain global in the sense that they can only capture the total amount of individual inbreeding. With genetic map information, one may alternatively rely on the identification of stretches of homozygous markers also referred to Runs of Homozygosity (RoH) (e.g., McQuillan *et al*, 2008) to estimate individual inbreeding at both a local genome scale and genome-wide (as the proportion of the genome contained in locally inbred regions). RoH are indeed most often interpreted as IBD chromosome segments that were inherited from a common ancestor without recombination (and mutation) in neither of them. Assessing the distribution of RoH within individual genomes has thus become popular to characterize inbreeding in a wide range of model species including humans (Kirin *et al*, 2010; McQuillan *et al*, 2008; Pemberton *et al*, 2012) or livestock (Bosse *et al*, 2012; Ferencakovic *et al*, 2013). RoH also allows to distinguish between recent and more ancient inbreeding (Kirin *et al*, 2010; Pemberton *et al*, 2012; Purfield *et al*, 2012) since pairs of IBD chromosomal segments tracing back to more remote ancestors are expected to be shorter because of a higher number of historical recombination events.

However, the main limitations of RoH–based approaches lie in their underlying rule-based procedure. For instance, the definition of the minimal number of homozygous markers (and segment length) and the maximum proportion of allowed heterozygous markers (to account for genotyping error) is mostly arbitrary. As a model-based alternative, Broman & Weber (1999) proposed a formal statistical approach to assess the IBD (or autozygous) status of the RoH they identified by accounting for population allele frequencies and genotyping error rates. Leutenegger *et al.* (2003) further provided a full probabilistic modeling of the IBD process along the chromosomes by developing a Hidden Markov Model (HMM). The HMM framework allows to make use efficiently of the available genetic information contained in the sequences of both homozygous and heterozygous markers and the linkage maps and can handle whole-genome sequence data (Narasimhan *et al*, 2016) including those obtained from low-fold sequencing experiments (Vieira *et al*, 2016). Although powerful, the aforementioned methods rely on a two-states HMM considering each marker either belongs to an IBD or a non-IBD chromosome segments. The transition probabilities between the (hidden) states of successive markers then depend on their given genetic distances, a parameter controlling the rate of changes per unit of genetic distance and the individual inbreeding coefficient. Considering only two states (IBD or non-IBD) thus amounts to assume that all the individual inbreeding originates from one or several ancestors in a single generation in the past and that all the IBD segments have the same expected length. However, in both natural and domesticated populations, the sources of individual inbreeding are multiple, since they are all related to their usually complex past demography history, making such an hypothesis of a single inbreeding event highly unrealistic.

We herein propose to extend previous HMM by considering several IBD-classes, each associated with a different inbreeding age. This new model allows to provide a better fit to individual genetic data (either genotyping or sequencing data) and to refine the genomic partitioning of inbreeding into stretches of IBD regions from possibly different ancestral origins. To evaluate the accuracy of the methods, we carried out comprehensive simulation studies. In addition, three real data sets from human, dog and sheep populations were analyzed in more detail to illustrate the range of application of the methods. As a by-product of this study, a freely available program, named ZooRoH was developed to implement inferences under the newly developed model.

## The Models

In the following we describe our HMM to model individual genomes as mixtures of IBD and non-IBD segments. We first consider a model with only two states (one IBD or autozygous class and one non-IBD class) and then describe the extension of the model to combine several IBD classes with varying time to the common ancestor (age measured in generations). To deal with the specificities of Next-Generation Sequencing (NGS) data (whole genome sequencing, low-fold sequencing, genotype-by-sequencing) that may provide less accurate genotype call than SNP chip arrays, we also propose alternative emission probabilities functions that integrate over the uncertainties of each possible genotype.

### The two–states model (1G model)

The 1G model is similar to the HMM previously proposed by Leutenegger *et al.* (2003) and assumes that the genome is partitioned in either IBD and non-IBD tracts that actually correspond to the two hidden states (*K* = 2). The 1G model further relies on a one order Markov process to define the transition probabilities between successive hidden states, such a modeling representing a good approximation of the IBD process along the chromosome in the absence of interference (Lander & Green, 1987; Leutenegger *et al*, 2003; Thompson, 2008). Consider two adjacent loci *M*_*l*−1_ and *M*_*l*_ separated by *r*_*l*_ Morgans (*l* > 1) and let *G* represent the size of the inbreeding loop i.e. twice the number of generations from a common ancestor and *ρ* the mixing coefficient corresponding to the proportion of IBD segments in the genome. Under the 1G model, *ρ* can be interpreted as a measure of the individual inbreeding coefficient *F* (Leutenegger *et al*, 2003). Let further *S*_*l*_ denote the (hidden) state of *M*_*l*_ with *S*_*l*_ = 1 and *S*_*l*_ = *K* = 2 for an IBD and non-IBD state respectively. The four transition probabilities between the hidden states of every pairs of consecutive markers are then defined as:

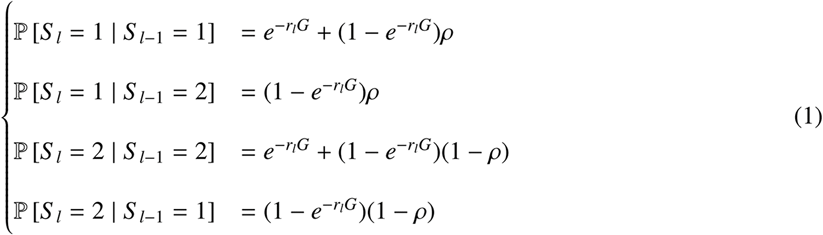

This amounts to assume that co-ancestry changes (leaving an IBD or non-IBD segment) between two adjacent markers *M*_*l*−1_ and *M*_*l*_ occur with a probability equal to 1 − *e*^*−rlG*^. It should thus be noticed that the same rate of co-ancestry changes (*G*) is used for both IBD and non-IBD tracks since we model the inheritance of chromosomal segments present in a single generation (that of the common ancestor). Under such assumptions, the length of IBD segments (inherited from a single ancestor) is exponentially distributed with an expected mean equal to 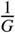. Because consecutive segments in the genome might belong to the same class, the overall lengths of the IBD and non-IBD segments have expected means equal to 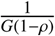 and 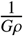 respectively (Leutenegger *et al*, 2003). Vieira *et al* (2016) also used a similar approach to model the transition probabilities whereas Narasimhan *et al* (2016) relied on a unique parameter for the transition probabilities that integrates both *G* and *ρ*.

– *l*

### Extension to multi-states models (KG models)

With a unique IBD class, the 1G model described above considers that all the IBD segments have approximately the same age either because they originate from a single ancestor (one strong inbreeding event) or from multiple ancestors in the same generation (e.g., during a bottleneck). Population history might however lead to far more complex patterns. For instance, common ancestors tracing back to different generations can be frequent in small populations, in populations under strong selection or in endangered populations with declining size. We therefore propose to extend the model to *K*_IBD_ different IBD classes, each characterized by their own mixing coefficient *ρ*_*c*_ and rate *G*_*c*_ (*c* ∈ (1, *K*_IBD_)). Note that *G*_*c*_ might be interpreted as twice the age (in generations) of the inbreeding class *c*. Common ancestors from IBD class *c* transmitted IBD segments whose lengths are exponentially distributed with a mean equal to 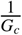. For the sake of generality, we may include several non-IBD classes but in the present study we only used one non-IBD class labeled *K* (i.e., the total number of classes *K* = *K*_IBD_ + 1) with a mixing proportion *ρ*_*K*_ and a change rate *G*_*K*_. The transition probabilities between the hidden states *S*_*l*−1_ and *S*_*l*_ of two adjacent loci *M*_*l*−1_ and *M*_*l*_ read:

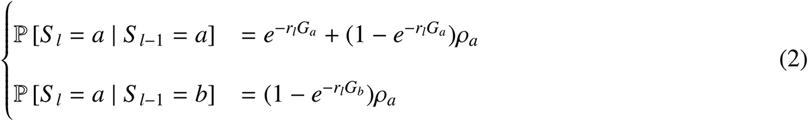

where *a* ∈ (1, *K*) and *b* ∈ (1, *K*) represents the identifier of the *K* different states (recalling that *K* also represents the non-IBD state). It is important to note that when *K* = 2, i.e. we only consider two states (*K*_IBD_ = 1 state and one non-IBD), the 2G model is slightly different than the 1G model since the two states are not constrained to have the same rate *G*.

### Emission probabilities and extension to NGS data

To complete the specification of the HMM we need to specify the emission probabilities, i.e., the probabilities of the data *Y*_*l*_ observed at each marker *M*_*l*_ given the underlying state *S*_*l*_ of the two individual chromosomes that might either be IBD (*S*_*l*_ ≠ *K*) or non-IBD (*S*_*l*_ = *K*). Let *I*_*l*_ represent the number of alleles observed for marker *M*_*l*_ (in the rest of the study we only considered bi-allelic SNP i.e., *I*_*l*_ = 2 for all *l*) and *A*_*li*_ the corresponding alleles (*i* ∈ (1, *I*_*l*_)). Depending on the technology and the analyses performed, *Y*_*l*_ then either consists of i) a genotype *A_li_A*_*lj*_ (where *i* ∈ (1, *I*_*l*_) and *j* ∈ (1, *I*_*l*_)) among the 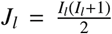 possible genotypes; or ii) a vector of likelihoods 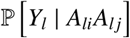 for each possible genotypes as provided by a genotype calling model as implemented within standard and popular softwares such as GATK (McKenna *et al*, 2010) or SAMTOOLS (Li *et al*, 2009). This allows to account for the genotype uncertainty which is highly recommended when dealing with NGS, particularly with low-fold sequencing data.

### Emission probabilities for genotyping data

Let *p*_*li*_ be the population allele frequency of allele *A*_*li*_ which is assumed to be known. If the two chromosomes are IBD in *M*_*l*_ (*S*_*l*_ ≠ *K*), we define the emission probabilities of the genotype *A_li_A*_*lj*_ as follows:

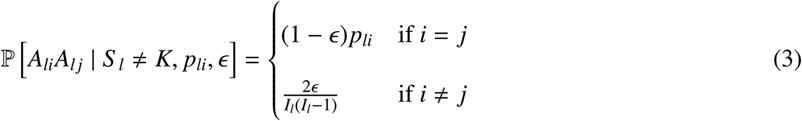

where *ϵ* is the probability (assumed to be known) to observe an heterozygous marker when the two underlying chromosomes are IBD in *M*_*l*_ either resulting from a genotyping error or a recent mutation. In other words, we assume that the vast majority of the polymorphic markers were segregating in the population before the common ancestors of the IBD segments and thus interpret recent mutations as genotyping errors. For non-IBD segments (tracing back to much more ancient ancestors), each genotype emission probabilities are derived assuming Hardy-Weinberg equilibrium and disregarding genotyping error (or recent mutation):

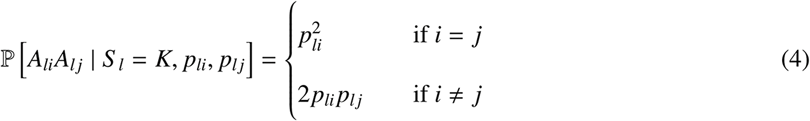

Note that these emission probabilities slightly differ from those considered in Leutenegger *et al.* (2003).

### Emission probabilities for genotype likelihood data

To account for genotype uncertainty, emission probabilities are obtained by integrating over all the possible genotypes:

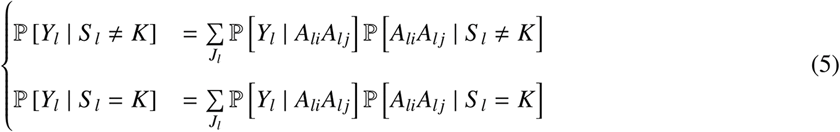

where 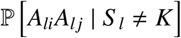 and 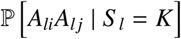 are as defined in equation 3 above (the error term *ϵ* then mostly capturing the effect of recent mutations). This modeling is similar to that recently proposed by Vieira *et al* (2016).

## Materials and Methods

### Inference

#### Estimation of model parameters

Assuming the population allele frequencies (*p*_*li*_) of each marker *M*_*l*_ and the error term *ϵ* are known, the set of parameters Θ that needs to be estimated consists of the mixing proportions *ρ* and the rates (interpreted as ages for the inbreeding classes) *G* of the defined IBD and non-IBD classes. Therefore, Θ consists of two parameters (*ρ* and one rate *G*) for the 1G model and 2*K* parameters for a multi-classes KG model (with *K*_IBD_ = *K* − 1 inbreeding classes). For multiple-IBD models, we alternatively consider reducing the parameter space by pre-defining the ages *G*_*k*_ of the *K* classes leading to only estimate the *K* mixing proportions *ρ*_*k*_ (hereafter called mixKG model). For all the models, parameter estimation was achieved with the Expectation-Maximization (EM) algorithm known as the Baum-Welch algorithm that is very popular in the HMM literature (Rabiner, 1989). The program ZooRoH implementing the algorithm for the different models is freely available at https://github.com/tdruet/ZooRoH. Unless otherwise stated, model parameters were estimated with 1000 iterations of the EM algorithm and setting *ϵ* to 0.001.

#### Estimation of the realized local (locus-specific) inbreeding (*φ*_l_)

The Baum-Welch algorithm allows to estimate the local state probabilities that correspond in our case to the *K* probabilities 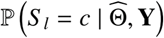 that the two chromosome segments belong to the IBD class *c* (*c* ∈ (1, *K*_IBD_)) or to the non-IBD class (*c* = *K*) at the marker *M*_*l*_ position given the estimated parameter set 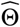 and the observed genetic data **Y**. These probabilities can be used to estimate both the realized genome-wide (over all the markers) and local (for each and every marker) inbreeding. Indeed, genetic data allows to directly infer the realized IBD status of an individual for each locus in the genome as opposed to pedigree-based inbreeding estimates that infer the expected IBD status for all the loci. More precisely, the local estimate 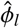 of the realized inbreeding at marker *M*_*l*_ is defined as the probability that this marker lies in an IBD segment and may thus be computed by summing over all its local IBD state probabilities (i.e., excluding the non-IBD class):

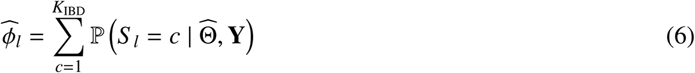

#### Estimation of the realized inbreeding associated to each IBD age-based classes 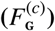 and the genome-wide inbreeding (*F*_G_)

As above, the inbreeding 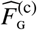 associated to IBD class *c* (*c* ∈ (1, *K*_IBD_)) can be defined as the proportion of the genome belonging to the class *c* and is estimated as the average of the corresponding local state probabilities over all the *L* locus:

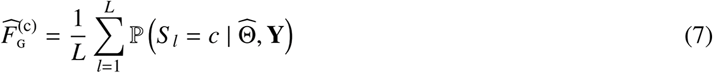

Finally, the genome-wide estimate of the realized individual inbreeding 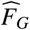 is simply the average over the genome of the local estimates obtained for the *L* markers:

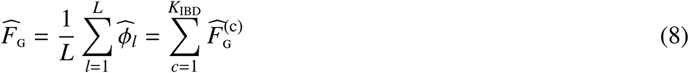

#### Model assessment

Because the optimal number of states (*K*_IBD_ or *K*) is usually unknown, we may be interested in characterizing, for a given data set, the strength of evidence for alternative number of states. To that end we relied on the Bayesian Information Criterion (**BIC**) which is a standard criterion for model selection among a finite set of models and was computed as:

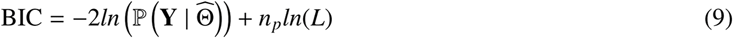

where 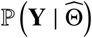 is the maximum of the likelihood function obtained with the estimated parameters 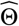 (computed with the forward algorithm (Rabiner, 1989)), *L* is the number of markers and *n*_*p*_ is the number of parameters, i.e., *n*_*p*_ = 2*K* − 1 for a KG model (with K-1 IBD classes) and *n*_*p*_ = *K* − 1 for a mixKG model (see above).

### Simulated data sets

#### Simulation under the inference model

The model was first tested by simulating data under the inference models. We simulated genotyping data at biallelic markers (SNPs) for 500 individuals considering a genome that consisted of 25 chromosomes of 100 cM length (i.e., 100 Mb length assuming a cM to Mb ratio of 1). The marker density was set to 10, 100 or 1,000 evenly spaced SNPs per Mb (i.e., 25,000, 250,000 or 2,500,000 SNPs in total). When simulating data under the 1G inference model, the individual genome is a mosaic of either IBD or non-IBD segments whose length is exponentially distributed with the same rate equals to the simulated *G* (twice the age in generations of the inbreeding event). For each chromosome in turn, we successively generated consecutive segments by sampling their length in the corresponding exponential distribution and randomly declaring them as IBD or non-IBD with a probability *ρ* and 1 − *ρ* (where *ρ* represents the simulated mixing coefficients). The process stops when the cumulative length of the simulated segments was greater than 100 cM (the last simulated segment being trimmed to obtain a chromosome length exactly equal to 100 cM). Under the multi-states model with several IBD classes, simulations were performed sequentially, with successive waves of inbreeding starting with the most ancient. We started by simulating the most ancient IBD class with the process described above. Then, each new IBD class was simulated similarly (with its own *G*_*i*_ and *ρ*_*i*_) except that new inbreeding (IBD) masked previous classes whereas non-IBD segments did not change previously simulated states.

To simulate genotyping data, we first randomly sampled for each SNP the population frequency of an arbitrarily chosen reference allele either i) from an empirical distribution derived from real cattle genotyping SNP assay and WGS data (Figure S1), or ii) from a (U-Shaped) distribution *β* (0.2, 0.2) that mimics NGS data (Figure S1). Given the simulated IBD status of the segments on which each SNP lie (see above), we used these sampled allele frequencies to simulate SNP genotypes as described for the emission probabilities above (eqs. 3 and 4). We used the parameter *ϵ* set to either 0.1% or 1% to introduce random genotyping errors (changing one genotype to one of the two other genotypes) and to evaluate the robustness of the models.

To simulate low-fold sequencing data (50 individuals) we sampled at each marker a number of reads *t* according to a Poisson distribution with mean *λ* (the average coverage). For homozygote genotypes (simulated as described above), the *t* sampled reads always carried the same allele (no sequencing error) and for heterozygotes, we used a binomial distributions (with parameters *t* and 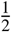) to sample the read counts for the two possible alleles. We then considered for each simulated SNP *l*, the read counts *t*_*l*1_ and *t*_*l*2_ observed for each of the two alleles to derive the three genotype likelihoods of the three genotypes *A*_*l*1_ *A*_*l*1_, *A*_*l*1_ *A*_*l*2_ and *A*_*l*2_ *A*_*l*2_:

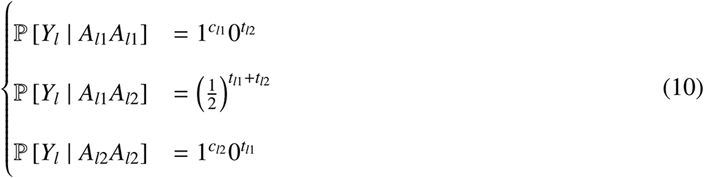

Finally, to assess the accuracies of the model estimation, we computed the Mean Absolute Error (MAE) for each parameter *α* of interest as:

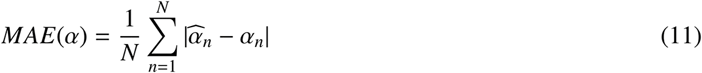

where *N* is the number of simulated individuals, 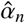 is the estimated parameter value for individual *n* and *α* is the corresponding simulated value.

#### Simulations under a discrete time Wright-Fischer process

The inference model we used is based on hypotheses (exponential distribution for length of IBD segments, Hardy-Weinberg equilibrium in non-IBD states, etc.) commonly used and that have been proven to work well (e.g., Leutenegger *et al*, 2003; Vieira *et al*, 2016). Still, we performed additional simulations relying on population genetics models to obtain simulated data less dependent on these assumptions. To that end we used the program ARGON (Palamara, 2016) that simulates data under a discrete time Wright-Fischer process.

With constant and large effective population size *N*_*e*_, inbreeding is expected to be low and to be spread over many generations. To concentrate inbreeding in specific age classes we simulated bottlenecks keeping large *N*_*e*_ outside these events to reduce the noise due to inbreeding coming from other generations. In the first scenario *WF*1, we considered an ancestral population *P*0 with a constant haploid effective population size equal to *N*_*e*0_=20,000 that split in two populations *P*_1_ and *P*_2_ at generation time *T*_*s*_ in the past with respective population sizes *N*_*e*1_=10,000 or 100,000 (according to the scenario) and *N*_*e*2_=10,000. During four generations centered around generation *T_b_* ≪ *T_s_* in the past, *P*_1_ experienced a bottleneck with an (haploid) effective population size equal to *N*_*eb*_ and recovered its initial size. Population *P*_2_ that always maintains a constant size is actually used to select markers that were also segregating in the ancestral population *P*_0_ (only markers segregating at MAF ≥ 0.05 in both populations *P*_1_ and *P*_2_ were kept for further analyses). The different simulation parameters are expected to have various impacts on the distribution of inbreeding. For instance for larger *T*_*s*_, inbreeding tends to accumulate after the two populations split and selected markers will have an older origin. Similarly, the larger *N*_*e*1_, the less inbreeding is accumulating outside the bottleneck while with smaller *N*_*eb*_, more inbreeding is created during the bottleneck. In total, 50 diploid individuals were simulated in both populations *P*_1_ and *P*_2_ considering a genome that consisted of a single chromosome of 250 cM length (i.e., 250 Mb assuming a cM to Mb ratio of 1). The mutation rate was set to *µ* = 10^−8^ and we use the functionalities of ARGON to identify all the IBD segments > 10 kb and to obtain their ages (generation time of the most recent common ancestor).

A second scenario *WF*2 was also considered for simulations in which similar parameters were used but the bottleneck occurred at generation *T*_*b*_ = 20 and *N*_*e*1_ was kept constant for subsequent and more recent generations (instead of returning to its initial size as in scenario *WF*1). This scenario with a strong reduction of *N*_*e*_ was aimed at mimicking livestock populations for which inbreeding is expected to be mostly due to ancestors in the most recent generations.

### Human, dog and sheep real data sets

For illustration purposes, we used publicly available genotyping data from *i)* the Human Genome Diversity Panel (HGDP) (Jakobsson *et al*, 2008) as downloaded from ftp://ftp.cephb.fr/hgdp_supp10/Harvard_HGDP-CEPH; *ii)* the dog LUPA project (Vaysse *et al*, 2011) as downloaded from http://dogs.genouest.org/SWEEP.dir/Supplemental.html; and *iii)* the Sheep Diversity panel (Kijas *et al*, 2012) as downloaded from the WIDDE database (Sempere *et al*, 2015). We then used the software plink (Purcell *et al*, 2007) to process and filter the genotyping data by removing individuals with a genotyping call rate below 90% and only keeping autosomal SNPs that had call rate > 95% and a MAF > 0.01 (in the original data set). As a result, the final data sets consisted of 304,406, 152,151 and 48,872 SNPs in human, dog and sheep respectively. For each species, we restricted our analysis to a subset of six populations corresponding to *i)* Karitiana (n=13), Pima (n=14), Melanesian (n=11), Papuan (n=17), French (n=28) and Yoruba (n=22) in humans; *ii)* Doberman Pinschers (n=25), Irish Wolfhounds (n=11), Jack Russell Terriers (n=12), English Bulldogs (n=13), Border Terriers (n=25) and Wolves (n=12) for the dog data set; and *iii)* Soay (n=110), Wiltshire (n=23), Dorset Horn (n=21), Milk Lacaune (n=103), Rasa Aragonesa (n=22) and Rambouillet (n=102) in sheep. Note that, within each population, markers with a MAF below 0.01 (within a population) were discarded from the analysis.

## Results

### Performance of the different models

#### Analyzing data simulated under the 1G inference model

We first analyzed individual genomes of 2,500 cM (with a marker density of 10 SNPs per cM) that were simulated under the 1G inference model, i.e., the simplest model. Depending on the two chosen simulation parameters (age of inbreeding *G* and mixing proportion *ρ*), these individual genomes thus consisted of a mosaic of IBD and non-IBD segments (in proportions *ρ* and 1 − *ρ* respectively) that both originated from the same ancestral generation (*G*/2 generations ago). In total, we analyzed with the 1G, the 2G, the 3G and the 4G models, 500 individuals per simulated scenarios, considering in total 33 different scenarios representatives of a wide range of values for both *G* (from *G* = 2 to *G* = 256) and *ρ* (from *ρ* = 0.0075 to *ρ* = 0.5). As mentioned in the Model section above, under the 1G model that was used for these simulations, *ρ* is highly similar to the individual inbreeding *F*_G_. The results obtained from the analyses under the 1G model are detailed in Table 1 for 20 different scenarios. In addition, tables S1 and S2 give the results from the analyses under all the four models (1G, 2G, 3G and 4G) for all the 33 different scenarios.

Overall, estimates of both model parameters 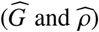 and individual inbreeding *F*_G_ obtained under the 1G model (Table 1 and Table S1) were found virtually unbiased and quite accurate (small MAE) irrespective of the considered scenarios. As expected, the 1G model performed even better when the number of IBD segments was higher and these were longer (smaller *G*) since more SNPs are available for their identification. For instance, for a given simulated 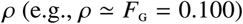, the MAE of 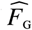 increased with larger simulated *G* (e.g., from 1.1 × 10^−3^ when *G* = 16 to 4.6 × 10^−3^ when *G* = 256). The performance of the 1G model to estimate local inbreeding (*φ*_*l*_) was further evaluated by computing the corresponding MAE either for all the SNPs 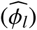 or for the SNPs lying within IBD segments only 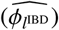 (Table 1 and Table S1). Note that for every simulated SNP *l*, the actual *φ*_*l*_ value is known (i.e., *φ*_*l*_ = 0 or *φ*_*l*_ = 1 if the SNPs is within a non-IBD or a IBD segment respectively). Hence, if the model performs well and all the *φ*_*l*_ are accurately estimated (i.e., 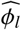 close to 0 or 1 for SNPs within a non-IBD or a IBD segment respectively), the MAE of 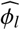 should be close to 0. Conversely, departure of the 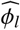 MAE from 0 indicates that IBD (respectively non-IBD) positions have non-zero probability to be non-IBD (respectively IBD). Besides, inspecting the 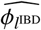 MAE allows to restrict attention to the prediction accuracy of truly IBD segments. As shown in Table 1, when inbreeding is recent (*G* < 32, i.e. less than 16 generations ago) MAE for both 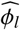 and 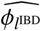 are close to 0 indicating that both IBD and non-IBD positions are correctly identified with a high support. Also, at constant level of overall (simulated) inbreeding (e.g., *ρ* ≃ *F*_G_ = 0.125) the accuracy decreases with higher value of *G* (e.g., from 1.0 × 10^−2^ when *G* = 4 to 2.1 × 10^−2^ when *G* = 8 for the 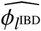 MAE). When considering more ancient (and/or) lower simulated inbreeding values, the 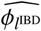 MAE increased faster than the overall 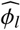 MAE. This indicates that there is not enough information (number of SNPs per IBD segments) to confidently classify some positions, in particular those within the shortest IBD segments, the longest IBS segments or the segments boundaries. It is however important to notice that the local inbreeding estimates 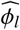 always remained very well calibrated, i.e., for any *p* ∈ (0, 1), the proportion of SNPs truly lying within IBD segments among the SNPs with 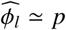 was close to *p* (Figure S2). Accordingly, and as mentioned above, the global estimators of individual inbreeding (*F*_G_) and the model parameters (*ρ* and *G*) remained accurate (Table 1).

**Table 1.**
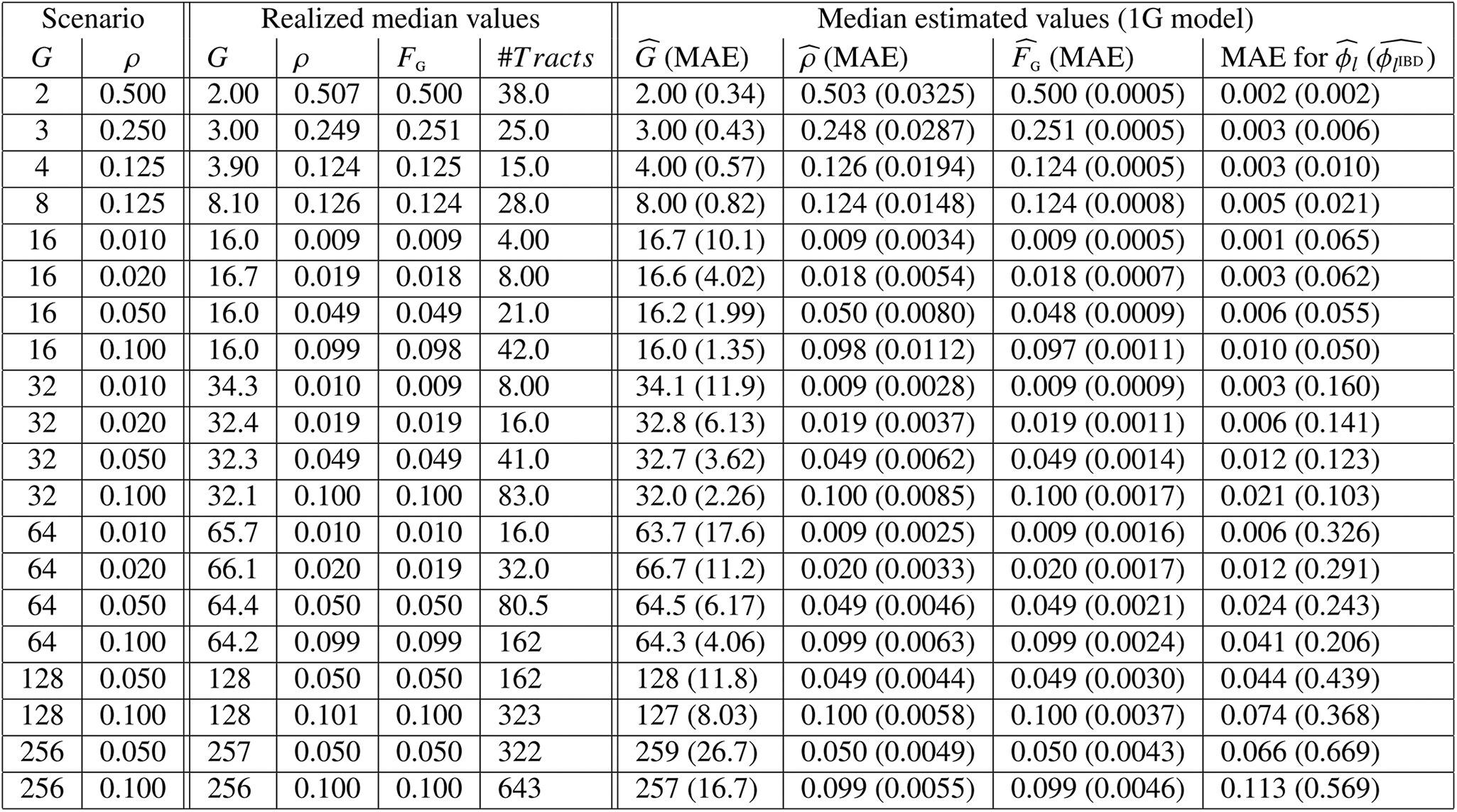
Performance of the 1G model on data simulated under the 1G inference model. The simulated genome consisted of 25 chromosomes of 100 cM with a marker density of 10 SNPs per cM. Genotyping data for 500 individuals were simulated under the 1G inference model for each of 20 different scenarios defined by the simulated *G* and *ρ* values reported in the first two columns. The table reports the resulting median realized (true) values (across the 500 simulated individuals) for the age of inbreeding (*G*), the mixing proportions (*ρ*), the individual inbreeding (*F*_G_) and the number of IBD tracks (#*Tracks*). Similarly, the table gives the median estimated values and the Mean Absolute Errors (MAE) for the age of inbreeding 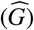, the mixing proportions 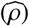 and the individual inbreeding 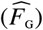. Finally, the table gives the MAE for the estimated local inbreeding (*φ*_*l*_) either for all the SNPs 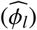 or for those actually lying within IBD segments 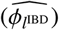.

| Scenario |  | Realized median values |  |  |  | Median estimated values (1G model) |  |  |  |
| --- | --- | --- | --- | --- | --- | --- | --- | --- | --- |
| $G$ | $\rho$ | $G$ | $\rho$ | $F_G$ | $\#Tracts$ | $\widehat{G}$ (MAE) | $\widehat{\rho}$ (MAE) | $\widehat{F}_G$ (MAE) | MAE for $\widehat{\phi}_I$ ($\widehat{\phi}_{IBD}$) |
| 2 | 0.500 | 2.00 | 0.507 | 0.500 | 38.0 | 2.00 (0.34) | 0.503 (0.0325) | 0.500 (0.0005) | 0.002 (0.002) |
| 3 | 0.250 | 3.00 | 0.249 | 0.251 | 25.0 | 3.00 (0.43) | 0.248 (0.0287) | 0.251 (0.0005) | 0.003 (0.006) |
| 4 | 0.125 | 3.90 | 0.124 | 0.125 | 15.0 | 4.00 (0.57) | 0.126 (0.0194) | 0.124 (0.0005) | 0.003 (0.010) |
| 8 | 0.125 | 8.10 | 0.126 | 0.124 | 28.0 | 8.00 (0.82) | 0.124 (0.0148) | 0.124 (0.0008) | 0.005 (0.021) |
| 16 | 0.010 | 16.0 | 0.009 | 0.009 | 4.00 | 16.7 (10.1) | 0.009 (0.0034) | 0.009 (0.0005) | 0.001 (0.065) |
| 16 | 0.020 | 16.7 | 0.019 | 0.018 | 8.00 | 16.6 (4.02) | 0.018 (0.0054) | 0.018 (0.0007) | 0.003 (0.062) |
| 16 | 0.050 | 16.0 | 0.049 | 0.049 | 21.0 | 16.2 (1.99) | 0.050 (0.0080) | 0.048 (0.0009) | 0.006 (0.055) |
| 16 | 0.100 | 16.0 | 0.099 | 0.098 | 42.0 | 16.0 (1.35) | 0.098 (0.0112) | 0.097 (0.0011) | 0.010 (0.050) |
| 32 | 0.010 | 34.3 | 0.010 | 0.009 | 8.00 | 34.1 (11.9) | 0.009 (0.0028) | 0.009 (0.0009) | 0.003 (0.160) |
| 32 | 0.020 | 32.4 | 0.019 | 0.019 | 16.0 | 32.8 (6.13) | 0.019 (0.0037) | 0.019 (0.0011) | 0.006 (0.141) |
| 32 | 0.050 | 32.3 | 0.049 | 0.049 | 41.0 | 32.7 (3.62) | 0.049 (0.0062) | 0.049 (0.0014) | 0.012 (0.123) |
| 32 | 0.100 | 32.1 | 0.100 | 0.100 | 83.0 | 32.0 (2.26) | 0.100 (0.0085) | 0.100 (0.0017) | 0.021 (0.103) |
| 64 | 0.010 | 65.7 | 0.010 | 0.010 | 16.0 | 63.7 (17.6) | 0.009 (0.0025) | 0.009 (0.0016) | 0.006 (0.326) |
| 64 | 0.020 | 66.1 | 0.020 | 0.019 | 32.0 | 66.7 (11.2) | 0.020 (0.0033) | 0.020 (0.0017) | 0.012 (0.291) |
| 64 | 0.050 | 64.4 | 0.050 | 0.050 | 80.5 | 64.5 (6.17) | 0.049 (0.0046) | 0.049 (0.0021) | 0.024 (0.243) |
| 64 | 0.100 | 64.2 | 0.099 | 0.099 | 162 | 64.3 (4.06) | 0.099 (0.0063) | 0.099 (0.0024) | 0.041 (0.206) |
| 128 | 0.050 | 128 | 0.050 | 0.050 | 162 | 128 (11.8) | 0.049 (0.0044) | 0.049 (0.0030) | 0.044 (0.439) |
| 128 | 0.100 | 128 | 0.101 | 0.100 | 323 | 127 (8.03) | 0.100 (0.0058) | 0.100 (0.0037) | 0.074 (0.368) |
| 256 | 0.050 | 257 | 0.050 | 0.050 | 322 | 259 (26.7) | 0.050 (0.0049) | 0.050 (0.0043) | 0.066 (0.669) |
| 256 | 0.100 | 256 | 0.100 | 0.100 | 643 | 257 (16.7) | 0.099 (0.0055) | 0.099 (0.0046) | 0.113 (0.569) |

As shown in Table S1, the estimates of *G* for the IBD class under the 2G model started to be substantially biased for scenario with *G* ≥ 128. More interestingly, the performances of the 2G model (Table S1) and both the 3G and 4G models (Table S2) were highly similar to those of the 1G model for the estimation of both genome-wide (*F*_G_) and local (*φ*_*l*_) individual inbreeding.

#### Analyzing simulated data with several underlying IBD classes

We further evaluated the performances of the different models on simulated data sets with more than one class for the underlying IBD segments, i.e. for which inbreeding originated from several sources of different ages *G*_*k*_ and contributions 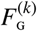 to the overall inbreeding. We detail hereafter the analyses of individual genomes of 2,500 cM (with a marker density of 10 SNPs per cM) that were simulated under the 3G inference model, i.e., assuming two different classes for IBD segments and one non-IBD class. Each simulation scenario was thus defined by the ages of inbreeding (*G*_1_ and *G*_2_) and the mixing proportions (*ρ*_1_ and *ρ*_2_) of the two classes of IBD segments. It should be noticed that the simulated mixing proportions (*ρ*_1_ and *ρ*_2_) directly controlled (and are generally close to) the amount of inbreeding originating from their corresponding IBD class. As shown in Table 2 for six different scenarios (and Tables S3 and S4 for a total of 23 different scenarios), estimates of the overall individual inbreeding (*F*_G_), of the ages (*G*_1_ and *G*_2_) and of the inbreeding contributions ( 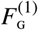 and 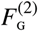) for the two IBD classes were close (but slightly biased) to the simulated values providing the differences between the ages of the two IBD classes was large enough (e.g., *G*_1_/*G*_2_ ≥ 16), i.e., the overlap between the distributions of the IBD segments lengths is reduced. As the difference between the ratio of successive *G*_*i*_ became smaller, all inbreeding tended to concentrate in the first IBD class that had an overestimated age for small simulated *G*_1_ (Table 2 and Table S3). For instance, for the scenario with *G*_1_ = 4 (*ρ*_1_ = 0.125) and *G*_2_ = 16 (*ρ*_1_ = 0.100), 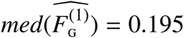 (*med* standing for median) and 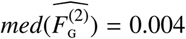 while 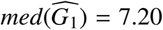 and 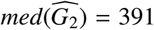 across the 500 simulated individuals (Table 2). Strikingly however, the overall individual inbreeding *F*_G_ always remained very well estimated with MAE≤ 0.005 for all scenarios (Table 2 and Table S4). Finally, as for the simulations under the 1G model previously considered, accuracy in the estimation of local inbreeding was found to mostly depend on the ages *G*_1_ and *G*_2_ (Table 2 and Table S5), the MAE for both 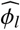 and 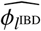 lying in a similar range than the one observed previously on data simulated under the 1G model. More precisely, given the relatively sparse SNP density considered, MAE remained accurate (i.e., ≤ 0.05) while *G*_1_ < *G*_2_ ≤ 64 but started to increase for higher values probably due to the inclusion of smaller IBD segments.

**Table 2.**
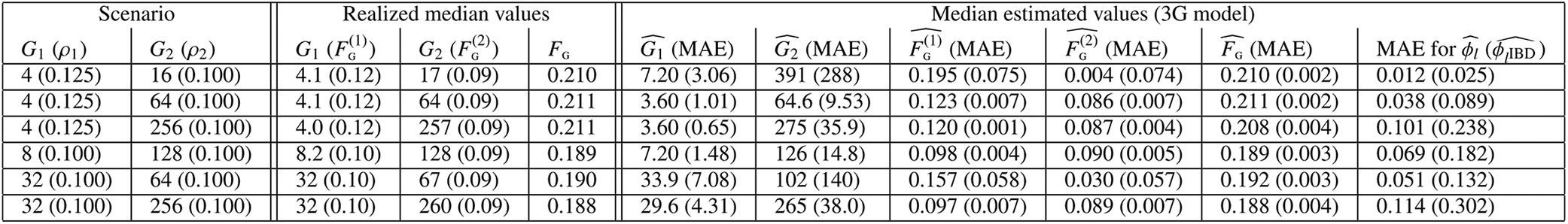
Performance of the 3G model on data simulated under the 3G inference model (i.e., two IBD classes and one non-IBD class). The simulated genome consisted of 25 chromosomes of 100 cM with a marker density of 10 SNPs per cM. Genotyping data for 500 individuals were simulated under the 3G inference model for each of 6 different scenarios defined by the simulated ages of inbreeding *G*_1_ and *G*_2_ (reported in the two first columns) and the corresponding mixing proportions *ρ*_1_ and *ρ*_2_ (reported in the third and fourth columns) of the two classes of IBD segments. The table reports the resulting median realized (true) values (across the 500 simulated individuals) for the ages of inbreeding (*G*_1_ and *G*_2_), the amount of inbreeding originating from each IBD class (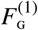 and 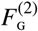) and the overall individual inbreeding (*F*_G_). The table further gives the median (and their associated MAE) of the estimated values 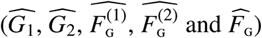 obtained under the 3G model. The table also gives the MAE for the estimated local inbreeding (*φ*_*l*_) either for all the SNPs 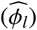 or for those actually lying within IBD segments only 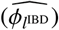.

| Scenario |  | Realized median values |  |  | Median estimated values (3G model) |  |  |  |  |  |
| --- | --- | --- | --- | --- | --- | --- | --- | --- | --- | --- |
| $G_1 (\rho_1)$ | $G_2 (\rho_2)$ | $G_1 (F_g^{(1)})$ | $G_2 (F_g^{(2)})$ | $F_g$ | $\widehat{G}_1$ (MAE) | $\widehat{G}_2$ (MAE) | $\widehat{F}_g^{(1)}$ (MAE) | $\widehat{F}_g^{(2)}$ (MAE) | $\widehat{F}_g$ (MAE) | MAE for $\widehat{\phi}_l$ ($\widehat{\phi}_{IBD}$) |
| 4 (0.125) | 16 (0.100) | 4.1 (0.12) | 17 (0.09) | 0.210 | 7.20 (3.06) | 391 (288) | 0.195 (0.075) | 0.004 (0.074) | 0.210 (0.002) | 0.012 (0.025) |
| 4 (0.125) | 64 (0.100) | 4.1 (0.12) | 64 (0.09) | 0.211 | 3.60 (1.01) | 64.6 (9.53) | 0.123 (0.007) | 0.086 (0.007) | 0.211 (0.002) | 0.038 (0.089) |
| 4 (0.125) | 256 (0.100) | 4.0 (0.12) | 257 (0.09) | 0.211 | 3.60 (0.65) | 275 (35.9) | 0.120 (0.001) | 0.087 (0.004) | 0.208 (0.004) | 0.101 (0.238) |
| 8 (0.100) | 128 (0.100) | 8.2 (0.10) | 128 (0.09) | 0.189 | 7.20 (1.48) | 126 (14.8) | 0.098 (0.004) | 0.090 (0.005) | 0.189 (0.003) | 0.069 (0.182) |
| 32 (0.100) | 64 (0.100) | 32 (0.10) | 67 (0.09) | 0.190 | 33.9 (7.08) | 102 (140) | 0.157 (0.058) | 0.030 (0.057) | 0.192 (0.003) | 0.051 (0.132) |
| 32 (0.100) | 256 (0.100) | 32 (0.10) | 260 (0.09) | 0.188 | 29.6 (4.31) | 265 (38.0) | 0.097 (0.007) | 0.089 (0.007) | 0.188 (0.004) | 0.114 (0.302) |

To provide insights on the behavior of our model to a misspecification of the underlying number of IBD classes, we also analyzed these data simulated under the 3G model with the 1G, the 2G and 4G models. As expected, when considering the 1G and 2G models, the estimated age of the single assumed IBD class was intermediate between the two simulated *G*_1_ and *G*_2_ actual values (Table S3). In agreement with previous findings, the 1G and 2G lead to highly similar estimates except for large *G*_1_ and *G*_2_ for which the estimated *G* tended to be higher with the 2G than the 1G model (e.g., 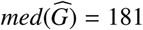 and 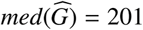 respectively for the scenario with *G*_1_ = 128 and *G*_2_ = 256). More interestingly, using the 1G and 2G models (i.e., with a single IBD class) to analyze these data resulted in an underestimation of *F*_G_ for scenarios with a marked differences between *G*_1_ and *G*_2_ (Table S4). Conversely, using an over-parameterized model such as the 4G did not introduce any additional bias compare to the 3G model. For instance, for the scenario with *G*_1_ = 4 (*ρ*_1_ = 0.125) and *G*_2_ = 256 (*ρ*_1_ = 0.100) that lead to a median realized inbreeding equal to 0.211 across the 500 simulated individuals, the median estimated inbreeding was equal to 0.162 with both the 1G and 2G models while it was equal to 0.208 and 0.209 with the 3G and 4G models respectively (Table S4). This suggested that the 1G and 2G model failed to capture some inbreeding. Accordingly, when focusing on the estimation of local inbreeding (Table S5), although the 1G and 2G models displayed a lower MAE for 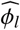 (i.e., computed over all the SNPs), this was essentially driven by SNPs lying in non-IBD segments. Indeed, both the 3G and 4G resulted in a lower MAE for 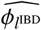 (i.e., computed over SNPs lying within IBD segments) suggesting these model allowed to better capture IBD segments at the expense of a slightly higher misassignment of SNP lying in non-IBD segments.

Overall, similar conclusions about the performance of the models to estimate the simulated parameters could be drawn when considering data sets with more than two underlying IBD classes (see Table S6 for results on data sets simulated and analyzed under the 4G model). It should however be noticed that increasing the number of IBD classes in the model also increased misassignment of IBD segments to their actual IBD class (Figure S3). In other words, some IBD segments, although correctly identified as IBD, might display a non-zero probability to belong to an incorrect IBD class (most generally a neighboring one). As a result, when increasing the number of simulated IBD classes, higher deviations of the estimated inbreeding age (*G*_*c*_) and contribution 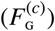 of each classes from their actual values could be observed (e.g., Table S6). Nevertheless, for higher ratio between successive class ages, these estimates remained fairly good. Importantly and as shown in previous simulations, the overall individual inbreeding (*F*_G_) was accurately estimated in all scenarios and MAE for local inbreeding mostly depended on the age of the IBD segments.

#### Using a set of *K* predefined IBD-classes (the mixKG model)

For a given model, instead of estimating the ages *G*_*k*_ of the different IBD classes, an alternative is to use a set of predefined age-classes and to only estimate the mixing proportions (*ρ*_*k*_). To illustrate and evaluate this strategy we hereby considered models consisting of 9, 11 or 13 IBD-classes depending on the simulated marker density (see below) and one non-IBD class leading to the so-called mix10G, mix12G and mix14G models according to our nomenclature. For each model, the predefined ages of the *K* − 1 IBD-classes always ranged from 2 to 2^*K*−1^ (with *G*_*k*_ = 2^*k*^ for each class *k* ∈ (1, *K* − 1)) while the age of the unique non-IBD class was the same as the older IBD class (i.e., *G*_*K*_ = *G*_*K*−1_ = 8192). Application of these mixKG models to the various data sets previously generated under the 1G, the 3G and the 4G inference models proved highly efficient (Table S7 and S8). For instance and in agreement with above results, the mix10G model provided accurate estimation of the overall inbreeding *F*_G_ (MAE always lower than 0.005 irrespective of the simulated scenarios) but also of the local inbreeding as indicated by MAE’s that were always as good as the best alternative model (e.g., compare Table S7 and Table S5). Moreover, such models with pre-defined ages for the IBD classes allowed to provide indications on the actual ages of inbreeding *G*_*k*_. We indeed observed that the estimated inbreeding contributions 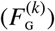 for the *K* − 1 IBD classes were mainly concentrated in those IBD-classes with pre-defined ages close to the true simulated ones as shown in Figure 1 for a dense SNP data sets (1000 SNPs per Mb) analyzed under the mix14G models and in Figures S4 to S8 for additional simulated data sets with smaller SNP density (either 10 or 100 SNPs per Mb) that were analyzed under mix10G or mix12G models.

**Figure 1.**
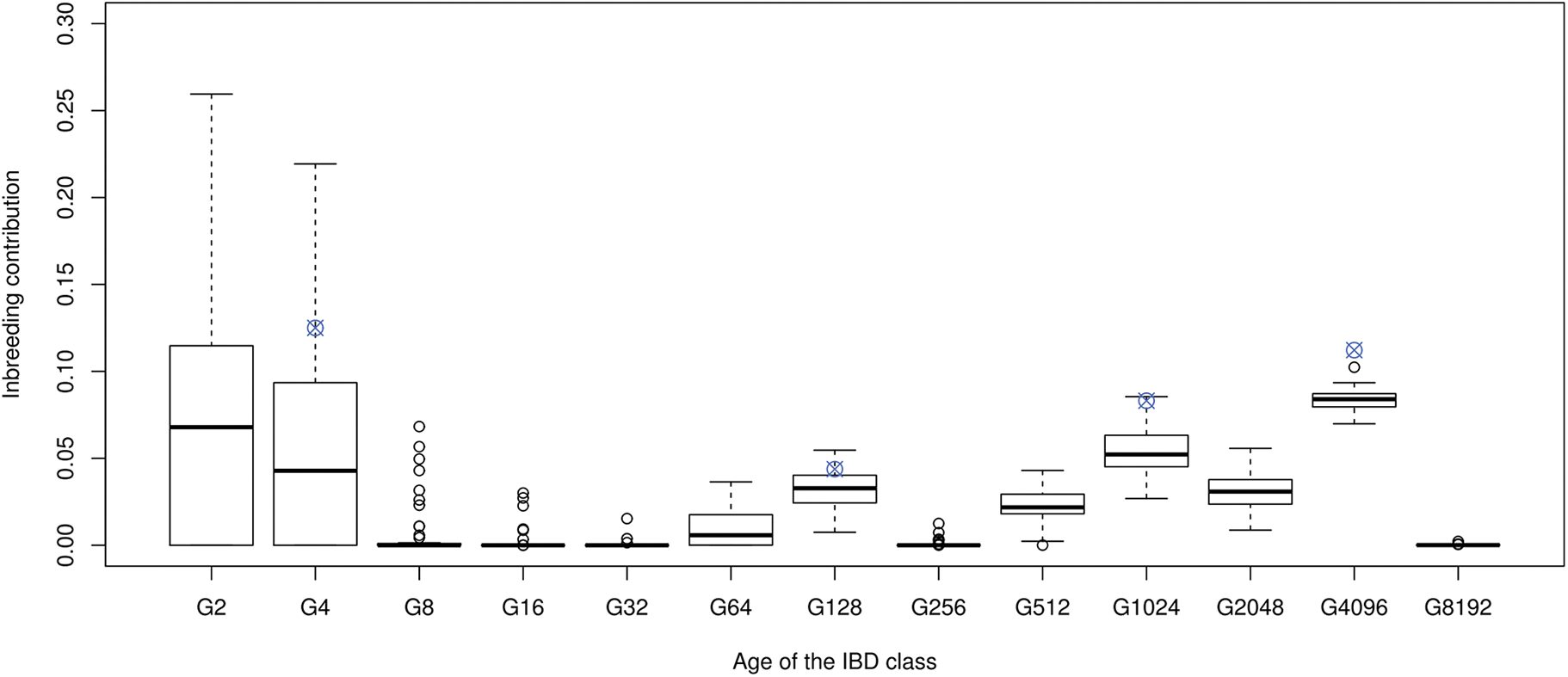
Estimated inbreeding contributions 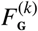 for 13 IBD classes with pre-defined ages (mix14G model) on data simulated under the 5G model (4 IBD classes). The simulated genome consisted of 25 chromosomes of 100 cM with a marker density of 1000 SNPs per cM. Genotyping data for 50 individuals were simulated under the 5G inference model i.e., with 4 IBD-classes with the following realized ages (inbreeding contributions) as indicated by a star in the plot: *G*_1_ = 4 ( 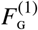 = 0.125), *G*_2_ = 128 ( 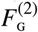 = 0.08), *G*3 = 1024 (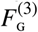 = 0.04) and *G*_1_ = 4 (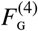 = 0.11). The data were analyzed with the mix14G that consisted of 13 IBD-classes with predefined ages ranging from 2 to 8192 (with *G_k_* = 2*^k^* for each class *k*) and one non-IBD class that had the same age as the older IBD class (i.e., *G*_*K*_ = *G*_*K*−1_ = 8192). For each of these 13 IBD classes, the boxplots give the distribution of the estimated inbreeding contribution 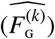 over the 50 simulated individuals.

#### Model comparisons and selection

We finally evaluated the BIC criteria to compare the models. When comparing different KG models (from 1G to 6G) applied to various simulation scenarios (ranging from 1 to 4 simulated IBD-distributions), we observed that the BIC criterion tended to support the correct underlying models and never provided support for models with a number of classes *K* higher than the simulated ones (Tables S9 and S10). Nevertheless, for simulations involving IBD segments from several classes (i.e., simulated under the 3G to 5G inference models), BIC may favor a model with a smaller number of IBD classes than the actual ones when the ages between successive classes are too close, although increasing SNP density improves the BIC resolution (Table S10). It should also be noticed that the BIC criterion never provided a stronger support in favor of the MixKG model (as defined above) when compared to the 6 others models considered (from 1G to 6G), possibly due to its higher number of parameters (e.g., *n*_*p*_ = 13 for the Mix14G model against *n*_*p*_ = 11 for the 6G model) (Tables S11 and S12). Yet, for simulations with several IBD classes (Table S12), the BIC support was generally higher than for the 1G and 2G models.

### Sensitivity of the models to genotyping error and marker informativeness

As only partially investigated above, when analyzing data with different SNP density, we expected that SNP information content, both in terms of marker density and genotyping accuracy, might be a key determinant of the resolution of the models. As a matter of expedience, we investigated this further by focusing on the 1G model (for both simulation and analyses) and evaluated the effect on its overall performances of changing the marker density and the SNP informativeness as summarized by the SNP allele frequency spectrum (AFS). Results confirmed that both the estimation of *G* and the identification of IBD positions associated to older inbreeding events always improved when increasing marker density and informativeness (Table 3). For instance, when the simulated *G* = 256, the MAE for 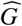 (respectively 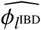) dropped from 36.9 (respectively 0.7313) with a marker density of 10 SNPs per cM and a *β* (0.2, 0.2) AFS to 8.06 (respectively 0.1994) with a marker density of 100 SNPs per cM and to 5.79 (respectively 0.0824) if, in addition, AFS was array-like. We also observe a better assignation of IBD segment to the correct IBD class with higher marker density (Figure S3). It is interesting to note that, at least for the range of parameters considered, *F*_G_ was accurately estimated irrespective of the marker densities and informativeness.

**Table 3.** Performance of the 1G model on simulated data sets with different SNP density and informativeness. The simulated genome consisted of 25 chromosomes of 100 cM with a marker density of either 10 or 100 SNPs per cM. Allele frequency spectrum (AFS) of each SNP reference allele were either sampled from an empirical distribution (array-like) derived from a real (cattle) genotyping assay (i.e., close to uniform) or from a (U-shape) *β* (0.2, 0.2) distribution that mimics NGS data. Genotyping data for 500 individuals were simulated under the 1G inference model for each of 3 different scenarios defined by the simulated *G* and *ρ* values reported in the first two columns. For each simulation, the table reports the resulting realized (true) median value (across the 500 simulated individuals) for the age of inbreeding (*G*) and the individual inbreeding (*F*_G_) together with the median of their estimated values 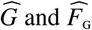 and corresponding Mean Absolute Errors (MAE). Finally, the table gives the MAE for the estimated local inbreeding (*φ*_*l*_) either for all the SNPs 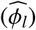 or for those actually lying within IBD segments only 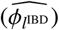.

| $G$ | $\rho$ | Simulation | | Realized median value | | Estimated median value | | |
| --- | --- | --- | --- | --- | --- | --- | --- | --- |
| | | SNP per cM | AFS | $G$ | $F_g$ | $\widehat{G}$ (MAE) | $\widehat{F}_g$ (MAE) | MAE for $\widehat{\phi}_l$ ($\widehat{\phi}_{IBD}$) |
| 4 | 0.125 | 10 | Array-like | 3.90 | 0.125 | 4.00 (0.57) | 0.124 (0.001) | 0.0026 (0.0101) |
| 4 | 0.125 | 100 | Array-like | 4.00 | 0.123 | 4.00 (0.51) | 0.123 (0.000) | 0.0002 (0.0009) |
| 4 | 0.125 | 10 | $\beta(0.2, 0.2)$ | 4.10 | 0.119 | 4.00 (0.64) | 0.120 (0.002) | 0.0068 (0.0272) |
| 4 | 0.125 | 100 | $\beta(0.2, 0.2)$ | 4.10 | 0.120 | 4.00 (0.55) | 0.120 (0.000) | 0.0006 (0.0023) |
| 64 | 0.100 | 10 | Array-like | 64.2 | 0.099 | 64.3 (4.06) | 0.099 (0.002) | 0.0410 (0.2056) |
| 64 | 0.100 | 100 | Array-like | 64.6 | 0.099 | 64.4 (2.00) | 0.099 (0.000) | 0.0035 (0.0181) |
| 64 | 0.100 | 10 | $\beta(0.2, 0.2)$ | 64.2 | 0.100 | 64.1 (6.26) | 0.100 (0.006) | 0.0807 (0.4032) |
| 64 | 0.100 | 100 | $\beta(0.2, 0.2)$ | 64.1 | 0.099 | 64.2 (2.50) | 0.099 (0.000) | 0.0095 (0.0482) |
| 256 | 0.100 | 10 | Array-like | 256 | 0.100 | 257 (16.7) | 0.099 (0.005) | 0.1134 (0.5689) |
| 256 | 0.100 | 100 | Array-like | 255 | 0.100 | 256 (5.79) | 0.100 (0.000) | 0.0164 (0.0824) |
| 256 | 0.100 | 10 | $\beta(0.2, 0.2)$ | 257 | 0.100 | 252 (36.9) | 0.100 (0.008) | 0.1462 (0.7313) |
| 256 | 0.100 | 100 | $\beta(0.2, 0.2)$ | 256 | 0.100 | 255 (8.06) | 0.100 (0.001) | 0.0398 (0.1994) |

We also investigated the sensitivity of the 1G model to the quality of genotyping or sequencing data. As shown in Table S13, when considering genotyping data (analyzed by setting *ϵ* = 0 for comparison purposes), we found that the presence of genotyping errors (either 1% or 0.1%) had little impact on the estimation of *F*_G_, moderate effects on the estimation of local inbreeding *φ*_*l*_ but estimates of *G* were strongly affected with an upward bias and an increased MAE. The magnitude of these effects was actually a function of the number of incorrect genotypes per IBD segment that increased the probability of observing heterozygotes and thus to cut the IBD segment into smaller ROH. As a result, the impact of genotyping errors was stronger for more recent inbreeding, at higher marker density and for higher error rate (Table S13). Interestingly, when analyzing the genotyping data with an appropriate error term i.e., setting *ϵ* = 0.01 (respectively *ϵ* = 0.001) for data simulated with a genotyping error of 1% (respectively 0.1%), the estimates of *G* became unbiased (Table S13). The accuracies with a 0.1% error were similar than without error but the MAE still remained larger with 1% genotyping errors and older inbreeding origins. Note that including a small error term in the model (*ϵ* ≠ 0) had little influence in the absence of genotyping errors.

We finally evaluated the sensitivity of the 1G model to various confidence levels in genotype calling by simulating data that mimic low-fold sequencing (or GBS) data for which several genotypes may have a non-zero probability. In these cases, read count data were simulated with a higher SNP density than above (1,000 SNP per cM) and variable coverage (from 1 to 10X). For each simulated SNP, the likelihood of the three possible genotypes were derived from the read count data as described in the Material and Methods section. The analyzed data sets then either consisted of i) the actual SNP genotypes (ideal situation) or ii) vectors of genotype likelihoods. As detailed in Table S14, we found that the model performed well in estimating the global parameters *G* and *F*_G_ with sequencing data. As expected, the performances improved with higher coverages and were similar than those obtained with the corresponding genotyping data as coverages ≥ 5X. Lowering sequencing coverages might indeed be viewed as decreasing SNP informativeness thereby leading to less accurate estimates for the different parameters (increased MAE), particularly for simulation in which inbreeding had an older origin (smaller IBD segments). For instance, for simulated *G* ≥ 512 and 1X coverage, both *F*_G_ and *G* were slightly underestimated (and to a lesser extent with 2X coverage) while for *G* ≤ 256, both global and local (*φ*_*l*_) estimates were accurate even with coverage as low as 1X (Table S14).

### Simulations under a discrete time Wright-Fischer process

To evaluate the robustness of the model to departure from model assumptions, we analyzed data simulated under a discrete-time Wright-Fisher process using the recently developed program ARGON (Palamara, 2016). For our purposes, a decisive advantage of ARGON is that it allowed to identify all the IBD segments (here we only considered those ≥ 10 kb) and to obtain their age (i.e., time to most recent ancestor or TMRCA). Inbreeding was generated by assuming population histories with either i) a strong bottleneck in the recent past followed by a rapid expansion as might be observed in invasive populations (WF1 scenarios) or ii) a reduced effective population size in the last twenty generations as might be observed in some domestic populations (WF2 scenarios). In total we considered 12 different WF1 scenarios and two different WF2 scenarios (see Material and Methods) and simulated 50 diploid individuals per scenario. As illustrated in Figure 2A for one WF1 scenario (see Figures S9 and S10 for all the 12 WP1 and the 2 WP2 scenarios respectively), the simulated history lead as expected to an enrichment in IBD segments that trace back to the bottleneck period within the simulated individual genomes (about 20% on average in Figure 2A). Yet, in most scenarios, a substantial proportion of inbreeding was associated to more ancient classes that accumulate inbreeding over many more generations. Indeed, a segment was considered IBD if it traced back to an ancestor from a generation more recent than the split time (*T*_*s*_ = 10^3^ or *T*_*s*_ = 10^4^ generations depending on the scenarios) of two modeled populations (see Material and Methods). Accordingly, in WF1 scenarios, this proportion increased with lower effective population size (*Ne*_1_), older split time (*T*_*s*_) and to a lesser extent higher bottleneck population size (*N*_*eb*_) and timing (*T*_*b*_) (Figures S9 and S10).

We analyzed all these simulated data sets with a mix10G model that consisted of 9 IBD-classes with predefined ages ranging from 2 to 512 (with *G*_*k*_ = 2^*k*^ for each class *k*) and one non-IBD class that had the same age as the older IBD class (i.e., *G*_10_ = *G*_9_ = 512). The choice for a MixKG model was motivated by our previous findings that demonstrated it was informative to date the origin of inbreeding and performed as well as other models in estimating local and overall inbreeding. In addition, it allowed to compare all the simulated individuals according to the same age-based partitioning of inbreeding.

**Figure 2.**
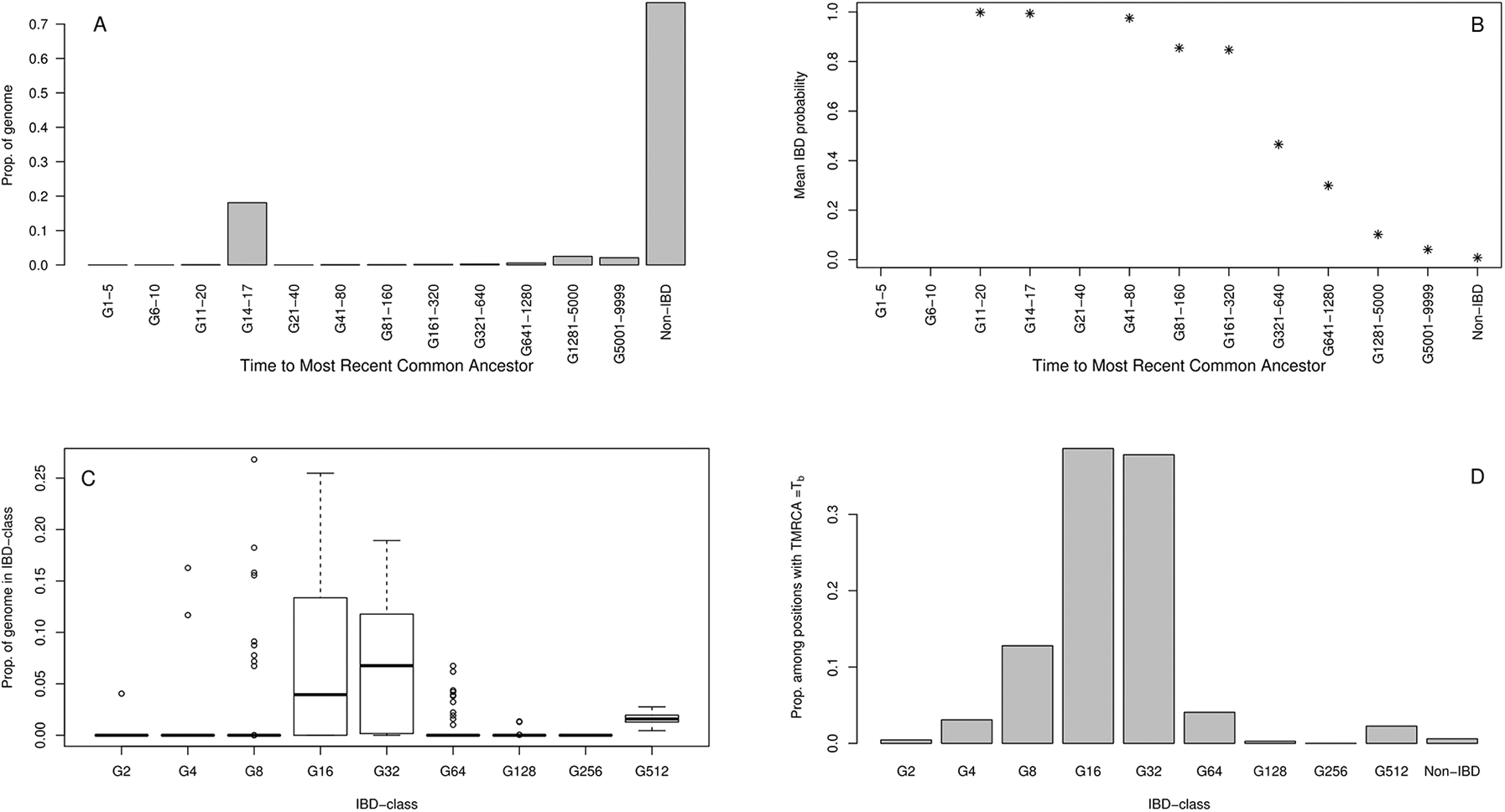
Evaluation of the mix10G model on a data set consisting of 50 diploid individuals simulated under a Wright–Fisher demographic history with varying population sizes. The population evolved under a WF1 scenario (see the Material and Methods section) with *Ne*_1_ = 10^5^, *T*_*s*_ = 10^4^ and a bottleneck lasting from generations 17 to 14 in the past and during which the population size was *N*_*eb*_ = 20. A) Realized distribution of the proportions of the simulated individual genomes lying within IBD segments as a function of their TMRCA (the interval G14-17 contains IBD segments tracing back to the bottleneck period, i.e., 14 to 17 generations backward in time) and within non-IBD segments (background). B) Estimated local inbreeding probabilities (*φ*_*l*_) averaged over all the simulated individuals and markers as a function of the actual TMRCA of the underlying IBD segments. C) Distributions of the estimated proportion of the individual genomes assigned to each of the 9 predefined IBD classes (over the 50 simulated individuals). D) Proportion of the SNPs lying in IBD segments originating from the bottleneck period (i.e., 14 to 17 generations backward in time) that are assigned to the 9 different IBD classes of the mix10G model (summed over all the 50 individuals).

As shown in Figure 2B (see Figures S11 and S12 for all the 12 WP1 and the 2 WP2 scenarios respectively), our HMM always allowed to efficiently identify IBD segments tracing back to common ancestors with TMRCA smaller than 80 generations, since the underlying SNPs displayed an estimated local inbreeding probability (*φ*_*l*_) close to one. In agreement with results obtained on simulations performed under the inference model (see above), the power to identify IBD segments of older origin gradually decreased (towards values almost always lower than 20% for TMRCA older than 5000 generations). Note that analyses of data sets simulated under the inference model showed that although the power was below one, overall inbreeding remained correctly estimated (see above). In addition, the model was found to perform well in assigning the identified IBD segments associated to the simulated bottlenecks since they were in their vast majority either assigned to their actual IBD class (i.e., with an age the closest to twice the age of the TMRCA) or to an immediately neighboring one. For instance, in the scenario considered in Figure 2, the estimated proportions of the individual genomes assigned to IBD segments were concentrated in the IBD class with predefined ages equal to 32 (*G*32), 16 (*G*16) and to a lesser (but less variable) extent in the oldest IBD-class (*G*512) (Figure 2C and Figures S13 and S14 for all the 12 WP1 and the 2 WP2 scenarios respectively). This was in agreement with the actual characteristics of the simulated individuals since IBD segments with a TMRCA≈ 16 that contributed on average to about 20% of their genome (Figure 2A) were mainly assigned (up to 70%) to the IBD classes *G*32 and *G*16 (Figure 2D). Note that the oldest IBD class G512 also captured some of these IBD segments together with a small proportion of those with an older TMRCA probably because these older IBD classes then become more frequent and have higher mixing coefficients. This effect was stronger when the bottleneck contributed less to the overall inbreeding and when the bottleneck was older. The performances of the model to correctly assign IBD segments however declined as the timing of the bottleneck was older or more generally as the proportion of inbreeding resulting from the period of reduced *N*_*e*_ was lower (Figures S13 and S14). Note that misassignment of IBD segments might also result from simulated segments being smaller/larger than expectations for a given pre-defined age *G*_*k*_ of the IBD class due to the stochastic nature of the Wright-Fisher process. In all cases however, we observed a peak of inbreeding in the IBD-class(es) corresponding to the period of reduced *N*_*e*_ or its neighbors (Figures S13 and S14). Overall, this simulation study thus confirmed that our model correctly identifies IBD-segments and gives good indications of the inbreeding’s age.

### Application to human, canine and ovine real data sets

We applied our model to individuals from human, dog and sheep populations, i.e., species representative of a wide range of demographic histories. Individuals were genotyped, as part of previous experiments (see Material and Methods) with assays containing various number SNPs (ca. 300K, 150K and 50K for human, dog and sheep individuals respectively) leading to different SNP density (ca., 1 SNP per 10kb, per 20 kb and per 60 kb respectively). As a result, and for the reasons mentioned above, the genotyping data were further analyzed with i) a mix14G model that consisted of 13 IBD-classes with predefined ages ranging from 2 to 8192 (with *G*_*k*_ = 2^*k*^ for each class *k*) and one non-IBD class that had the same age as the older IBD class (i.e., *G*_14_ = *G*_13_ = 8192) for humans and dogs; and ii) a mix9G model that consisted of 8 IBD-classes with predefined ages ranging from 2 to 256 (with *G*_*k*_ = 2^*k*^ for each class *k*) and one non-IBD class (*G*_10_ = *G*_9_ = 512) for sheep to account for the smaller SNP density. To interpret the results, it is useful to remind that the ages *G*_*k*_ of the predefined classes are approximately twice the TMRCA and that populations have variable ratio between genetic and physical distances when averaged between sexes: 1.16 cM/Mb for human (Kong *et al*, 2010), 1.26 cM/Mb for sheep (Johnston *et al*, 2016) and 0.88 cM/Mb for dog (Campbell *et al*, 2016). Indeed, we used for the analyses the SNP position on the physical maps accompanying the respective data sets. The estimated contribution of each pre-defined IBD class (averaged over all the individuals) are detailed for each populations and each species in Figure 3.

**Figure 3.**
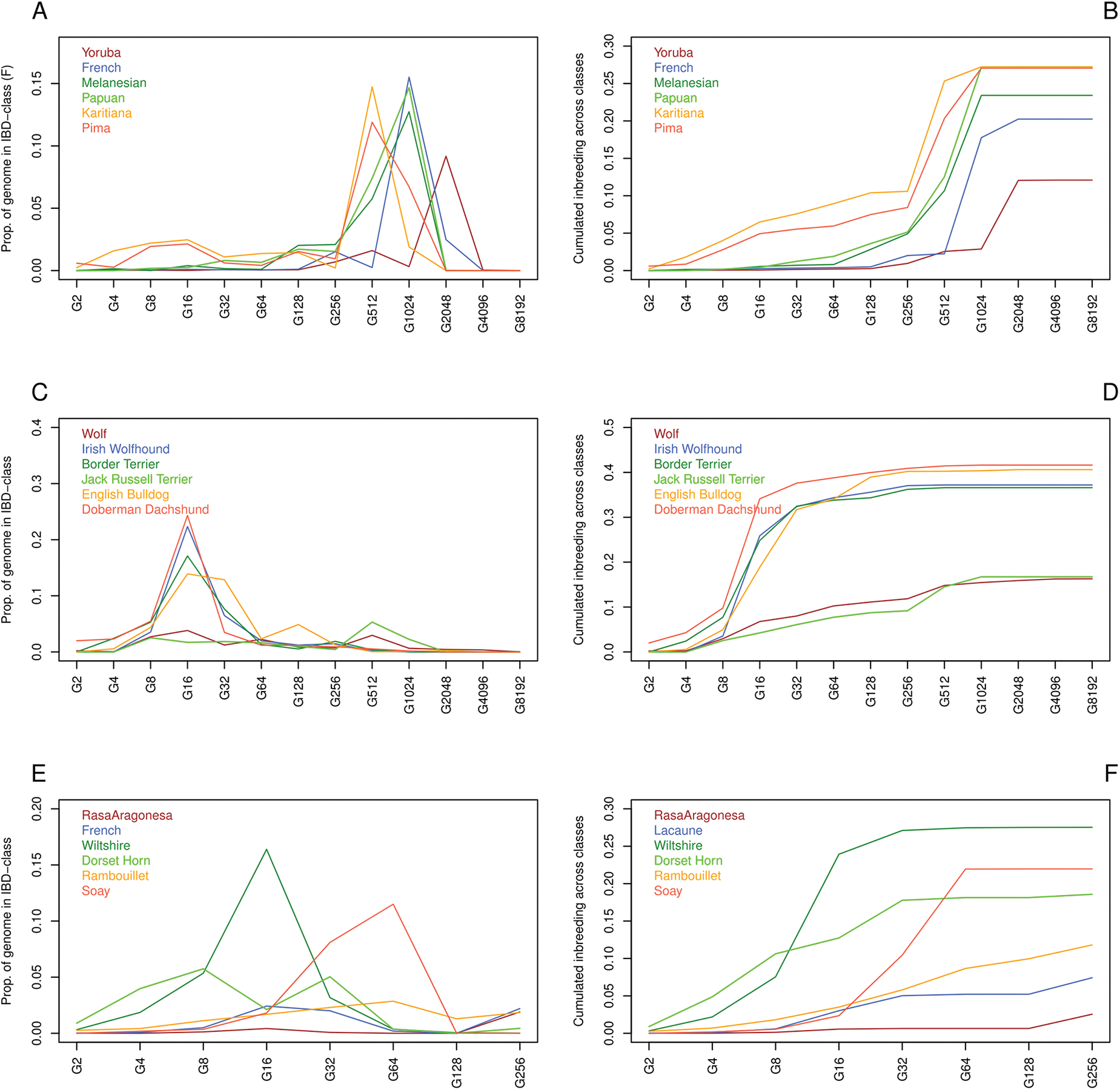
Average estimated proportions of inbreeding contribution of a set of *K* predefined IBD classes for human (A, *K* = 13), dog (C, *K* = 13) and sheep (E, *K* = 8) populations and corresponding average cumulative inbreeding (B, D and F for human, dog and sheep populations respectively).

Regarding humans, the six populations considered here (French, Yoruba, Melanesian, Papuan, Pima and Karitiana) have already been thoroughly analyzed in other studies (e.g., Jakobsson *et al*, 2008) including a study that provided a detailed assessment of the distribution of ROH of different lenghts (Kirin *et al*, 2010). Our results showed that the amount of overall inbreeding increased from Africans, Europeans, Oceanians to Native Americans from Central and Southern America with a generally remote origin (Figure 3A,B and Figure S17). More precisely, the ages of the main contributing IBD-classes that were generally consistent within population were clearly related to the *N*_*e*_ of the corresponding populations (the older the larger). Hence, the peak of inbreeding was i) in the class with *G*_*k*_ = 512 for Pima and Karitianas; ii) in classes with *G*_*k*_ = 512 and *G*_*k*_ = 1024 for Papuans and Melanesians; in the class with *G*_*k*_ = 1024 for French; and iv) in the class with *G*_*k*_ = 2048 for Yoruba. Interpretation of such old inbreeding, more related to population characteristics than individual differences, must be done with caution (see Discussion). Nevertheless it should be noticed that in French or Oceanian populations we observed some individuals with more recent inbreeding but this remained limited compared to Pima and Karitiana where there is strong evidence of recent inbreeding, some of the individuals having more than 10% inbreeding in very young classes from *G*_*k*_ = 2 to *G*_*k*_ = 8 (Figure 4A and Figure S17). These observations are consistent with previous findings by Kirin *et al* (2010) based on ROH that suggested the presence of both recent (long ROH) and ancient (short ROH) inbreeding in Native Americans. Conversely, individuals from Oceanian populations did not display long ROH (several Mb long) but had an excess of ROH of intermediate length (between 1 and 2 Mb) indicating a reduced *N*_*e*_ in the past. Finally, individuals from European and African populations mostly showed background inbreeding (short ROH) that correlated with the underlying *N*_*e*_. One major difference of our results with the aforementioned study by Kirin *et al* (2010) is that they only considered ROH > 500 kb leading to a lower estimated value (most probably downwardly biased) for the overall individual inbreeding.

Modern dog breeds present large amounts of inbreeding and are known to have experienced strong bottlenecks associated with the recent breed creation from a small number of founders (e.g., Vaysse *et al*, 2011). In addition, strong artificial selection and matings in small closed populations further contributed to increase inbreeding in the last decades (Lewis *et al*, 2015). Accordingly, as shown in Figure 3C,D and Figure S18, we observed massive inbreeding (sometimes higher than 20%) in the IBD-class with *G*_*k*_ = 16 (a common ancestor approximately 8 generations ago) in all the five breeds we analyzed but the Jack Russell Terrier that has a larger *N*_*e*_ (Vaysse *et al*, 2011). As expected also, wolves that did not experienced domestication did not present such an excess of inbreeding in recent generations. In each population (including wolves), some individuals were found to be highly inbred with an *F*_G_ ≈ 50% and approximately 25% of this inbreeding associated to an estimated common ancestor living only one or two generations ago (Figure 4B and Figure S18).

Finally, among the six sheep populations we investigated, three (the Rasa Aragonesa, Milk Lacaune and Rambouillet) displayed a large *N*_*e*_ (> 700) as described in Kijas *et al* (2012). Hence, individuals from the Rasa Aragonesa displayed almost no trace of inbreeding (max = 1.3% when cumulated up to the IBD-class with *G*_*k*_ = 8) while the cumulative inbreeding remained lower than 5% on average for individuals from the Milk Lacaune and Rambouillet breeds up to classes *G*_*k*_ = 32 (Figure 3E,F and Figure S19). Yet, some Rambouillet individuals presented high levels (> 20%) of recent inbreeding (Figure 4C and Figure S19). Conversely, the Wiltshire (*N*_*e*_ = 100) and Dorsethorn (*N*_*e*_ = 137) populations that went through a strong reduction in size in the early 1900’s (Dorsethorn to a lesser extent) were both found to have a high level of recent inbreeding (Figure 3 and Figure S19). The main contributing IBD-class was the one with age *G*_*k*_ = 16 for Wiltshire and *G*_*k*_ = 4 to *G*_*k*_ = 32 for Dorsethorn. Interestingly, the Wiltshire individuals were sampled from a New-Zealand flock that experienced several strong and successive bottlenecks in its recent history. Indeed, its founders were imported in 1974 from Australia where the breed had previously been introduced in 1952 and survived as a remnant population of as few as 12 ewes (O’Connell *et al*, 2012). Assuming a generation time of approximately 4 years in sheep, the distribution of the contribution of the most recent classes to the overall inbreeding is thus consistent with this demographic history. The sixth sheep population we investigated was the well known Soay sheep that had an estimated *N*_*e*_ = 194 (Kijas *et al*, 2012) and experienced a strong founder effect since the current population derives from a flock of 107 individuals that were transferred on the Hirta island in 1932 and then lived in complete isolation (Clutton-Brock & Pemberton, 2004). We observed for this population a small amount of recent inbreeding (for IBD classes with age *G*_*k*_ ≤ 16), even lower than in Milk Lacaune or Rambouillet, but rather high levels of inbreeding associated with IBD classes of ages between between 32 and 64 generations (Figure 3E,F and Figure S19). Integrating over all the generations, the Soay sheep thus appeared on average even more inbred than Dorsethorn, which explains the small estimated *N*_*e*_. However, despite this strong founder effect and the high resulting inbreeding level, we observed almost no individual with an inbreeding *F*_G_ > 5% in the most recent generations. The Soay breed represents an interesting example of a wild population resulting from a founder effect and in expansion. To summarize, our model allowed to provide deeper insights into the very different patterns of individual inbreeding observable in the sheep breeds. Indeed, these inbreeding patterns ranged from small as in the Rasa Aragonesa or limited level (with a few overly and recently inbred individuals) as in the Rambouillet breed, to moderate to high inbreeding level that either originated from strong bottleneck in the very recent (Wiltshire) or recent (Soay) past, or that resulted from the cumulative effect of a less pronounced population size reduction over more generations (Dorsethorn).

**Figure 4.**
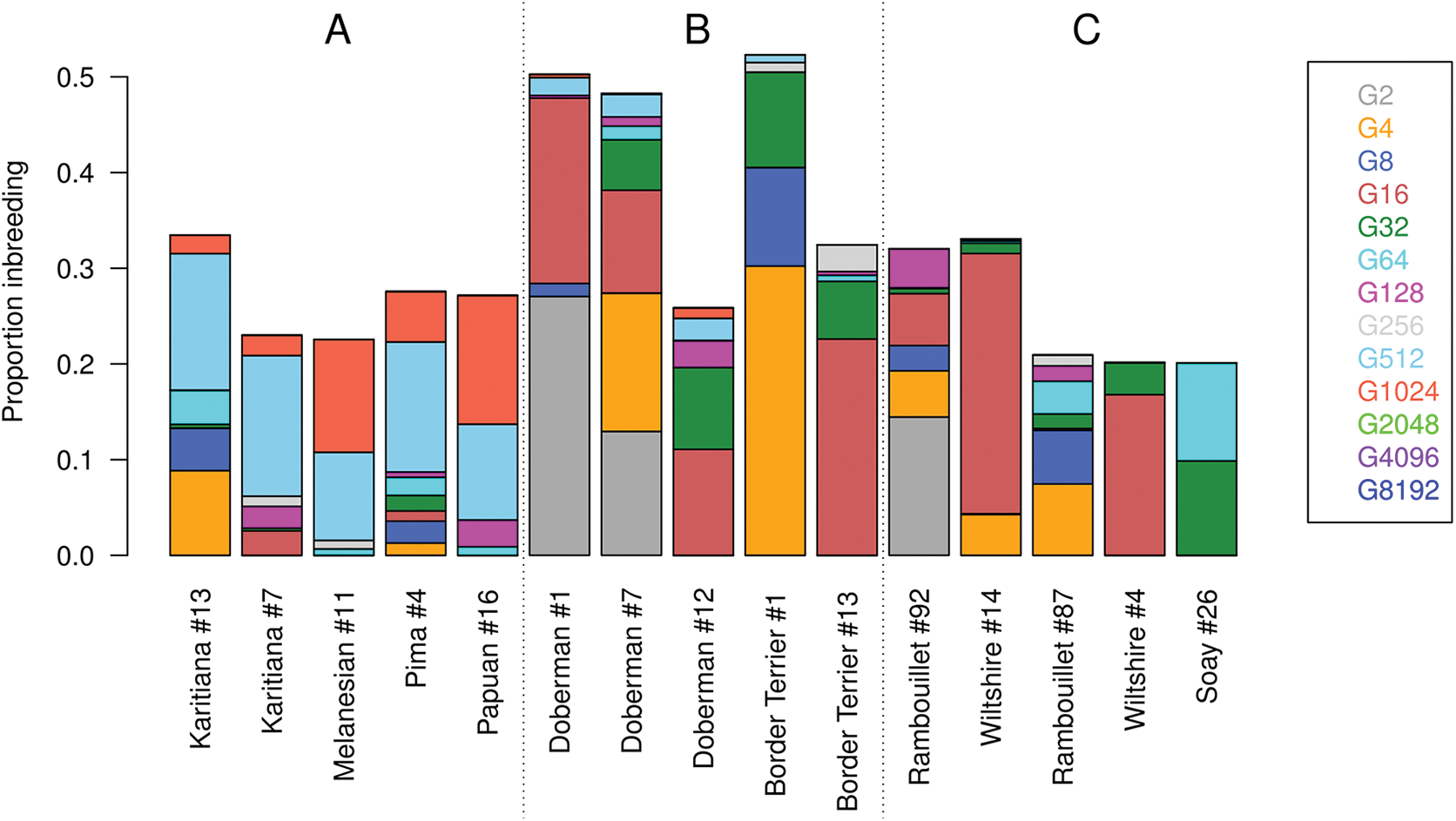
Estimated partitioning of inbreeding in five humans (A), five dogs (B) and five sheeps (C).

Importantly, besides providing a global estimator of inbreeding for each individual, the model also informs on the partitioning of this individual inbreeding which is highly valuable. For instance, individuals born from extremely consanguineous marriages might be easily identified. As an illustration, Figure 4B showed three dogs (Doberman #1 and #7, Border Terrier #1) that displayed approximately 25% inbreeding associated with the *G_k_* = 2 or *G_k_* = 4 IBD-class (ancestors living one or two generations ago) unlike other dogs from the same population (Doberman #12 and Border Terrier #13). These three individuals are likely resulting from matings between a sire and its daughter. This indicates that inbreeding is still present in these populations and is not only due to the breed creation event but to further management practices. High level of inbreeding associated to parents or grand-parents are also observed in sheep (19.2% for Rambouillet #92 in Figure 4C) and even in human (8.9% for Karitiana #13 in Figure 4A). For all these individuals, however, these recent events accounts only for a fraction of total inbreeding and a substantial proportion of inbreeding is due to more remote ancestors. More generally, by partitioning the total amount of inbreeding among ancestors from different generations, our model provides a better understanding of the origins of inbreeding in each individual. Hence, individuals with a similar overall inbreeding might display a quite different pattern of ancestral contributions captured by our model. For instance, for the three sheep individuals (Rambouillet #87, Wiltshire #4 and Soay #26) represented in Figure 4C that all displayed an overall inbreeding of approximately 20%, the inbreeding is mostly associated to the IBD-class *G*_*k*_ = 16 for the Wiltshire #4, to the two IBD-classes *G*_*k*_ = 32 and *G*_*k*_ = 64 for the Soay #26 whereas for the Rambouillet #87 individual, ancestors contributing to inbreeding trace back to a wide spectrum of generations (from *G*_*k*_ = 4 to *G*_*k*_ = 256). These observations are consistent with patterns at the population level. Interestingly, individuals with higher levels of inbreeding (Wiltshire #14 and Rambouillet #92) display comparable patterns with inbreeding concentrated in the IBD-class *G*_*k*_ = 16 for Wiltshire #14 and associated to several IBD classes for Rambouillet #92 (Figure 4C). In humans (Figure 4A), Native Americans from Central and Southern America were found to display different make-ups than Oceanians with similar levels of overall inbreeding (e.g., Karitiana #7 vs Melanesian #11 or Pima #4 vs Papuan #16). As expected from previous results, Oceanians actually displayed little traces of very recent inbreeding but accumulated more inbreeding in distant generations.

## Discussion

In this study, we developed and evaluated HMM models that use genomic data to estimate and to partition individual inbreeding into classes of different ages. There actually exist a wide variety of methods to estimate individual inbreeding and these have different properties. Pedigree-based methods rely on a genealogy (the inbreeding can only result from individuals within the genealogy) and predict the expected IBD status at a locus whereas genomic measures estimate realized inbreeding (the observed level of inbreeding). Genomic estimates can either be global, giving a unique measure per individual, or local. Obviously, these latter measures provide more information but require a higher marker density. Assessing the distribution of ROH within individual genome have recently become popular to characterize global and local inbreeding (Kirin *et al*, 2010; McQuillan *et al*, 2008; Pemberton *et al*, 2012). Most often, however, estimators relying on ROH are categorizing pairs of chromosome segments as IBD or non-IBD and do not provide intermediate values. They rely on the assumption that if stretches of homozygous markers are sufficiently long, they are IBD. Many parameters must be defined (including minimal number of homozygous markers, minimal length of an homozygous track, maximal spacing between successive markers, maximal number of heterozygous SNPs in a RoH) and these depend on the population under study and on the genotyping technology used. HMM’s as those developed in this study make a better use of all the information since they take into account the marker allele frequencies, the genotyping error rates, the genetic marker map (the genetic distance between successive markers) and the expected length of IBD tracks. Initially designed for genotyping arrays (Leutenegger *et al*, 2003), they can easily be extended to NGS data (Narasimhan *et al*, 2016) including low-fold sequencing data (Vieira *et al*, 2016) or genotype-by-sequencing data as done in our study, whereas simple ROH are inappropriate in such conditions. HMM’s also allow to automatically estimate some parameter of interest such as the frequency of IBD segments (a measure similar to the expected inbreeding if only one IBD-class is modeled) and their expected length. Finally, when relying on the Forward-Backward algorithm (as in our study), these models integrate all the available information to estimate the IBD probabilities of each marker in opposition to a binary classification as obtained with ROH or with a Viterbi algorithm in HMM (Leutenegger *et al*, 2003; Narasimhan *et al*, 2016; Vieira *et al*, 2016). Using a probabilistic model is particularly valuable when information is sparser and classification is more uncertain (e.g., for smaller and older IBD tracts, at lower marker density or informativeness, with higher genotyping error rates or with low-fold sequencing).

The most simple HMM we considered consists of a single IBD state (1G model) and is similar to several previously proposed ones (Leutenegger *et al*, 2003; Narasimhan *et al*, 2016; Vieira *et al*, 2016). This amounts to either assume that a single common ancestor is responsible for inbreeding or that the vast majority of IBD segments trace back to ancestors that lived in the same past generation. However, most populations have complex demographic histories, with varying *N*_*e*_ and common ancestors of IBD segments are thus expected to originate from many different generations in the past. As shown by our application in real data sets, even in domestic populations for which inbreeding might be expected to result from a limited number of founder individuals, individual inbreeding generally result from ancestors in different generations back in time probably due to the subsequent intense use of some key (selected) breeders. Hence, extending the model to several IBD-classes is highly valuable and might be viewed as defining multiple reference populations instead of a single one. Inbreeding is then captured as distantly in the past as made possible by the available marker density and informativeness. There is thus no need to arbitrarily define any base population with unrelated ancestors nor to select an arbitrary threshold below which stretches of homozygous markers are considered non-IBD. The first benefit of a multiple IBD-classes model is to better fit the data and to obtain more accurate estimators of inbreeding both locally and globally. Indeed, our simulations under the inference model with several IBD classes clearly showed that the 1G (and 2G) model underestimated *F*_G_ as some IBD segments were missed while the power to detect IBD segments was decreased. In addition, in the presence of ancient inbreeding, 1G model will tend to interpret recent (and thus longer) IBD segments as consecutive smaller segments of older origins because the estimated age of the single IBD class would tend to be older. Of course, in the absence of genotyping errors, the entire segment would then be correctly declared IBD and would appear as a long tract. However, at higher genotyping error rates (as with NGS data) such segments would be cut into smaller pieces. This would not happen when analyzing data with a model with multiple classes since recent IBD segments would then be associated to a class with a smaller age and the penalty in the HMM to leave the IBD-class and start a new IBD segment would be too large. Under a single-IBD class model, the age of the longest ROH further tends to be overestimated which might introduce substantial biases in applications that rely on the age of the IBD tracts to estimate some parameters of interest (e.g., the mutation rate). With two states HMM (Leutenegger et al., 2003), LD pruning is sometimes used to get rid of background LD and to force the model to concentrate on recent inbreeding (and hence avoiding the aforementioned problem). With multiple IBD-classes model (> 2*G* models), ancient inbreeding associated with background population LD is automatically assigned to the eldest IBD classes making LD pruning unnecessary for that purpose. Also, HMM with multiple IBD classes allows to determine whether there is a single or multiple IBD distribution(s) and to obtain information on the relatively recent demographic history of the population, providing *N*_*e*_ was reduced at some recent time in the past such as in populations under conservation or invasive populations. Such a modeling actually explores more recent generations and can be considered as complementary to approaches that infer past *N*_*e*_ thousands generations ago and many more as proposed by Li & Durbin (2011). Application to real populations demonstrated than the model can capture very different patterns including presence or absence of consanguineous matings, large *N*_*e*_ and low inbreeding, bottlenecks at varying time in the past, founder effects and reduced *N*_*e*_ due to isolation in the past (*G_i_* ≥ 100). Finally, with multiple classes, we can clearly identify individuals from extreme consanguineous matings (sire x daughter, first cousins, etc) because the recent inbreeding due to this recent ancestor is distinguished from the background inbreeding. Such examples with 25% inbreeding in class *G_i_* ≤ 4 were observed in dogs or sheep populations.

Our modeling approach actually allows to explore inbreeding in several dimensions: the global (*F*_G_), the local (*φ*_*l*_) and age-variable 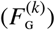. It has been stated that more ancient inbreeding should not be considered since deleterious variants are expected to be rapidly purged from populations. Yet, the number of generations for this purging to complete depends on the population history. For instance, strong bottlenecks tend to reduce the efficiency of purging deleterious variants (”The cost of domestication”) and artificial selection might favor some breeders carrying deleterious variants. Thanks to our model we could estimate the inbreeding depression associated with different age-classes. This requires appropriate data sets (individuals genotyped at high marker density to capture old inbreeding and with own fitness records) and sufficient variation in all IBD-classes. Alternatively, recent and old inbreeding can be compared by functional annotations of different segments. For instance, Szpiech *et al.* (2013) showed that long ROH are enriched for deleterious variants in humans. We can also use our model to test for local inbreeding depression and identify regions or variants where homozygosity seems more deleterious (e.g., Leutenegger *et al*, 2006).

Several strategies can be used to infer inbreeding in populations with our model. First, when using only one IBD class as in Leutenegger *et al* (2003), we can either estimate a single age common to both IBD and non-IBD classes or a different value for both states. The first option results in a model similar to Leutenegger *et al* (2003) and Vieira *et al* (2016) (note that the model by Narasimhan *et al* (2016) does not estimate the age but a single transition parameters combining *G* and the mixing proportions) and results in better estimates of age. Next, we can select the best number of IBD-classes according to the BIC criterion to compare the different models. When evaluated under simulated data, the BIC appeared to be conservative since the selected values were smaller or equal to the simulated ones. Note that with this approach we select the number of classes that best fit the data (merging several close classes if necessary) and not the real number of classes. Finally, we can use a set of IBD (and non-IBD) classes with predefined ages (the so-called MixKG models). It is then recommended to well separate these ages (e.g., using a ratio of 2 between successive ages to limit the overlap between the exponential distributions assumed for the IBD segment lengths) and cover a range of generations compatible with the available marker density. That strategy proved particularly efficient in most cases since it provided accurate estimates of the overall and local inbreeding while providing insights into the partitioning of inbreeding in the different age-classes and more easily comparable results across individuals from the same population. Such a model was only sub-optimal when a single and rare IBD class was simulated (which might not be usual in real populations) but required larger computational resources since more classes are simultaneously fitted.

Some precautions must be taken regarding interpretation of results. In our model, the mixing proportion and the rate of the exponential distribution are both estimated contrary to the model by Narasimhan *et al* (2016) where a single parameter is used. The mixing proportion can be interpreted as an expected inbreeding (the proportion of IBD segments among all segments) only if we have one IBD class and a single estimator for the age *G*. When segments from different distributions have different lengths, that interpretation is no longer correct (see the model section). The estimation of *G* might further be influenced by approximations in the model since we assume that the map is known without error, the recombination rate is not variable, there is no mutation and the population allele frequencies are known and did not vary over time. The estimation of this parameter is based on the distribution of lengths of IBD segments but this is a random process, for the same true *G* we can obtain segments of different lengths. For estimation from few IBD segments, the relative variation is higher. The presence of multiple-IBD classes generates also noise and the estimated distributions are often combinations of true underlying distributions. Therefore, the estimated inbreeding distribution must not be considered as exact but rather indicative. This is particularly true for ancient inbreeding classes for which there is less information and approximations are accumulated over many generations. Ancient inbreeding captures ancient demographic history (past *N*_*e*_ and resulting LD) and presents less variation among individuals (ancient inbreeding is the results of many lineages and variance decreases for large samples). Note that some methods do not consider as inbreeding such shorts ROH reflecting homozygosity for ancient haplotypes and contributing to local LD patterns although Broman & Weber (1999) declared that homozygosity resulting of “linkage disequilibrium is indeed the result of the mating of (very distantly) related individuals”. With our model, such inbreeding is automatically associated to ancient IBD-classes and separated from more recent inbreeding. Hence, users are free to interpret this as true inbreeding or background LD and to carry out LD pruning prior to analysis. The purpose of our model is to estimate individual genomic inbreeding and we advice to interpret only classes presenting enough variation. For older events and population parameters, we recommend to rely on other complementary models. Globally, estimation of different parameters is less accurate when less information is available (fewer IBD segments and less informative marker per segment). The model relies on two important hypotheses. First, it is assumed that most of the variants trace back further in time than the ancestors: the mutation did not happen in the path between the individual and its ancestor. With standard mutation and recombination rates (e.g., as in human or cattle), few mutations per IBD segment are expected on these paths (the value is relatively constant regardless of the age since older segments are smaller but have more time for mutations). So, as long as enough SNPs are present per segment, the impact of mutations should be low and accounted for by the genotyping error rate parameter. In addition, markers from genotyping arrays are old due ascertainment bias favoring polymorphism in several populations. Still the model should be used cautiously when this condition is not met e.g., in populations distantly related from all of those represented in the discovery panel of the genotyping array (monomorphic SNPs in all the individuals of the considered population should then be discarded). The second hypothesis is that the marker allele frequencies in the base populations are known but we have only estimates. A special attention must be taken when working with several very different populations and markers have been selected based on their frequencies in one of these. When many markers are not segregating in one population (due to ascertainment bias) but frequencies are estimated across populations, these markers will be considered variable. Their fixation in the breed might then be considered as inbreeding. It is therefore important either to estimate the frequencies within population or use markers segregating in all the populations.

We are working on several extensions of our model, for instance to better take into account the possibility of mutations or to estimate the allele frequencies. Another possible extension to capitalize on individual inbreeding for past demographic inference of the whole population would be to explicitly relate the contribution of each IBD-class to each and every individual inbreeding to the corresponding past effective population size and further consider all the individuals jointly to estimate these (hyper–)parameters. Such a development might be viewed as an extension of our individual-oriented model to the population level.

## Acknowledgements

We thank the Human Genome Diversity Project, the LUPA consortium and the International Sheep Genomics consortium for data sharing. John McEwan helped us to obtain the sheep data set and shared his knowledge on history of different sheep populations. The ZooROH project and this work were supported by the Fonds de la Recherche Scientifique FNRS (F.R.S.-FNRS) under Grant J.0134.16. Tom Druet is Research Associate from the F.R.S.-FNRS. We used the supercomputing facilities of the “Consortium d’Equipements en Calcul Intensif en Fédération Wallonie-Bruxelles” (CECI), funded by the F.R.S.-F.N.R.S.

## Data Accessibility

All data sets used in the present study are publicly available. the Human Genome Diversity Panel (HGDP) data was downloaded from ftp://ftp.cephb.fr/hgdp_supp10/Harvard_HGDP-CEPH, the dog LUPA project from http://dogs.genouest.org/SWEEP.dir/Supplemental.html and the Sheep Diversity panel from the WIDDE data base (Sempere *et al*, 2015). The program ZooRoH implementing our model can be freely obtained at https://github.com/tdruet/ZooRoH.

